# Optimal spatial release strategies for confined gene drives and *Wolbachia*

**DOI:** 10.64898/2026.03.04.709515

**Authors:** Ziye Wang, Jackson Champer

## Abstract

Gene drives are genetic elements that can rapidly spread through populations, offering potential solutions for controlling disease vectors and pests. In some scenarios, it is necessary to utilize drives that can be confined to only target populations. The success of these threshold-dependent gene drives, which require a minimum local frequency to establish, depends critically on the spatial strategy used for introduction. Here, we use a reaction-diffusion model to systematically identify optimal release patterns that maximize the per-capita efficiency for four distinct gene drive designs as well as use of *Wolbachia* bacteria, which spread similarly to frequency-dependent gene drives. We find that the most efficient release strategy is highly dynamic, transitioning from a broad “everywhere” release for short timeframes to a “multiple-ring” pattern for intermediate times, and finally to a focused “center” release for longer timeframes. These timeframes depend on the specific type of drive, with more powerful variants transitioning more quickly to center releases. Our results demonstrate that these optimized, variable release strategies can be substantially more effective than simple uniform releases. This study provides a quantitative framework for designing effective gene drive implementations, highlighting that a carefully planned spatial strategy is essential for maximizing impact, making optimal use of available resources.

## Introduction

Gene drives are genetic elements that can bias their inheritance and spread through populations [1–4]. This property makes them powerful potential tools for addressing major public health and ecological challenges. Modification drives can modify vector populations to prevent disease transmission, and suppression drives can suppress target pest species [5–7].

Gene drive systems can be broadly categorized based on their invasion dynamics into non-threshold and threshold drives [4, 8, 9]. The introduction threshold is the critical frequency above which the drive has to be released to spread successfully in a panmictic population. Zero-threshold drives, such as most CRISPR homing drives [10, 11], can invade a population from an arbitrarily low initial frequency. In contrast, threshold-based drives have a nonzero minimum introduction frequency, which can arise both from the intrinsic property of the drive mechanism itself or from any fitness cost imposed by the drive construct. Underdominance drives are a major type of confined drives that have a nonzero introduction threshold even in ideal form [9, 12–22]. The principle behind this is that drive/wild-type individuals are less fit than either wild-type or drive homozygotes, though the mechanism of this can vary between drives. Many additional drives lack an introduction threshold in ideal form, but gain one if they have any fitness cost (most real world drives are expected to have at least a small fitness cost due to the presence of the drive itself, regardless of drive mechanism). Examples of gene drives that have a threshold only with a fitness cost include *Medea* [23] and TARE [24, 25]/ClvR [26, 27], the latter of which belong to the family of CRISPR-based toxin–antidote drives. Only some of these drives can be used for population suppression, but all could potentially be used as part of a tethered drive system, allowing them to provide confinement for a homing suppression drive [22, 25, 28–30]. Maternally inherited *Wolbachia* has similar properties, though it usually has a substantial fitness cost and thus in practice would usually have population dynamics closer to underdominance systems [31, 32]. It has the benefit of often being able to block transmission of certain diseases on its own, unlike gene drives that require a cargo gene. It has thus had some success in the field [31, 32]. On the other hand, it is also less flexible due to lack of methods for engineering its genome.

The existence of an introduction threshold provides a highly desirable form of biocontainment, as the drive is less likely to invade non-target populations when migration rates are sufficiently low. On the other hand, it presents a significant implementation challenge: a release that is too small will fail, with the drive being rapidly eliminated by selection. This can be easily understood in a panmictic population, but in a spatial population, the dynamics of the drive are more complicated in situations where the threshold is exceeded in only some of the total area.

The theoretical basis for the spatial spread of such systems was established early, describing the advance of an alleles as a traveling wave [33]. Analysis of the dynamics of systems with a frequency threshold showed that the wave front of a threshold-based drive is “pushed” by dispersal from high-frequency areas behind it, in contrast to the “pulled” wave of a non-threshold drive [34]. This makes the spread of confined drives highly sensitive to spatial factors, such as density gradients and dispersal barriers, which can halt or even reverse their advance [35, 36]. Consequently, the spatial pattern of the initial release is not merely a logistical detail but an essential determinant of whether the drive can establish a self-sustaining wave of advance, which is possible only if the release exceeds a “critical bubble” [34, 36, 37]. Thus, some modeling work has investigated the release pattern for different systems, often aiming to find the least number of drive individuals required for success of a gene drive [16, 38–40]. However, such a “critical” gene drive release is not the optimal release strategy. It provides only the minimum required release for the drive to persist, not to ensure optimal spread. A true optimal release strategy would maximize the per-capita efficiency of the drive individuals released, defined as the final number of drive alleles for a given release effort, given a particular timeframe, These previous studies have shown significant variation in drive spread efficiency with different release patterns. Therefore, choosing the optimal release strategy can save considerable resources that can be deployed elsewhere.

In this study, we systematically identify the release patterns that maximize drive spread efficiency for four threshold-dependent gene drive systems and *Wolbachia*. Using a reaction-diffusion model, we find that the optimal release strategy is highly dynamic, transitioning from a broad “everywhere” release for short timeframes to a multiple-ring pattern for intermediate times, and finally to a focused single-region release for longer timeframes. Our results provide a quantitative framework for designing release strategies and demonstrate that an optimized spatial configuration can dramatically increase the efficiency of confined population genetic engineering systems.

## Methods

### Overview

To identify release strategies that maximize efficiency, we assessed the population dynamics of four gene drives plus *Wolbachia* bacteria with a reaction-diffusion model. These were selected to include both modification and suppression drives with a range of introduction threshold frequencies. We defined a “spread efficiency” metric based on the number of drive alleles generated (for modification drives) or population size reduction (for suppression drives) per released individual over a fixed time period. We then used numerical optimization to find the radially symmetric release pattern that maximized this efficiency for a given operational timeframe.

### Drive types

**Toxin-Antidote Recessive Embryo (TARE)** consists of a drive element inserted into a haplosufficient but essential gene [24, 25, 29]. The drive’s Cas9 and gRNA constitute the toxin and cleave the wild-type allele of the gene, while a recoded, cleavage-resistant rescue is provided in the drive itself. In germline cells, drive allele will cleave wild-type alleles, forming disrupted alleles, which are then inherited by offspring. If an individual has a drive mother, any wild-type alleles can be similarly cleaved at the early embryo stage. Disrupted alleles are recessive lethal, meaning that homozygotes for this allele are nonviable (Figure 1A). Any individual with at least one drive or wild-type allele is viable. Hence, TARE drive increases in frequency by eliminating wild-type alleles, which are turned into disrupted alleles and lost in disrupted homozygotes. Previous modeling studies found that TARE drive has an introduction threshold of zero, but gains a nonzero threshold whenever the drive allele has any fitness costs [24, 25, 41]. In this study, we assume an ideal TARE drive. It has no fitness costs for drive alleles and 100% germline and embryo cut rate.

**Figure 1.**
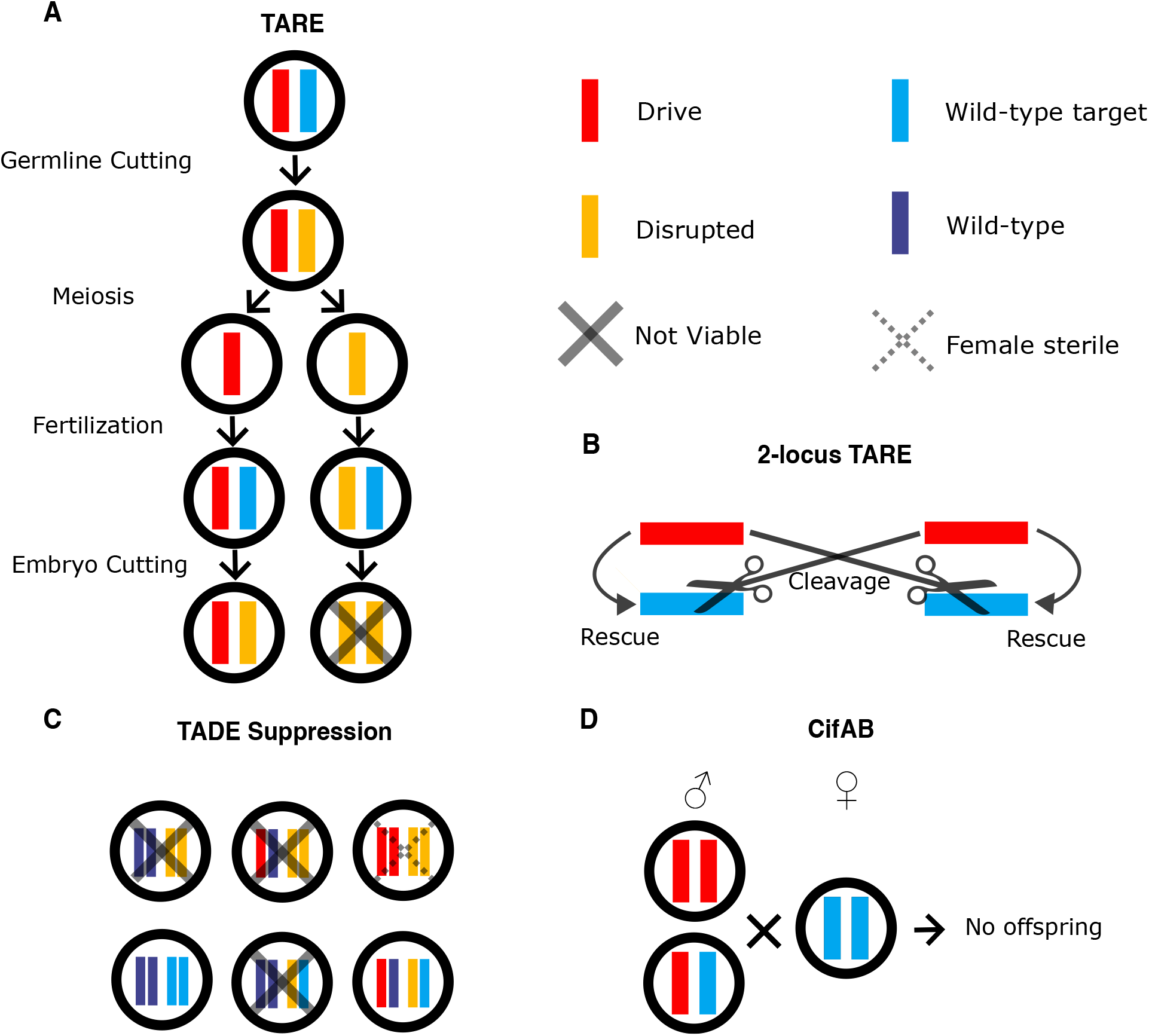
Schematic of gene drive mechanisms. **A TARE**. In the germline, the drive cleaves wild-type alleles, converting them to disrupted allele. In the progeny of drive females, maternally deposited Cas9 and gRNA will further cleave wild-type alleles. Disrupted allele homozygotes are nonviable. **B 2-locus TARE**. There are two TARE drive, and each cleaves the wild-type allele that the other rescues. **C TADE suppression**. Individuals with less than two copies of the target gene (either in the drive or wild-type allele) are nonviable. The drive construct is inserted into and disrupts a haplosufficient but essential female fertility gene. This makes female drive homozygotes sterile. **D *Wolbachia* CifAB**. A wild-type female mating with a male containing at least one copy of the drive allele will produce no offspring. Other types of mating have normal inheritance. For *Wolbachia*, wild-type females produce no offspring when mating with infected males, and *Wolbachia* infection is inherited maternally.

**2-locus TARE** is a drive that comprises two types of TARE constructs, each at different unlinked genetic loci [22]. Each drive allele provides rescue for the gene at its own site and cleaves the wild-type allele at the site of the other drive allele type (Figure 1B). Unlike TARE drive, 2-locus TARE has a moderate introduction threshold of 18% even without fitness costs for the drive alleles [22, 41]. We model ideal 2-locus TARE drive with no fitness costs and 100% germline and embryo cut rate.

**Toxin-Antidote Dominant Embryo (TADE) suppression** targets a haplolethal gene, where two functional copies are required for viability [25]. A recoded rescue of the target gene is also contained in the drive construct. Thus, if the number of drive and wild-type alleles in a genotype is less than two, the organism will be nonviable. TADE modification drive does not differ in any other respects from TARE drive in terms of its design. Unlike TARE drive, embryo cleavage is not desired in TADE because it would render some drive-carrying individuals nonviable. This would create a nonzero introduction threshold, even if all other performance aspects are ideal. A TADE suppression specifically disrupts a haplosufficient but essential female fertility gene, making females sterile if they lack a wild-type copy of the gene [22, 25, 41–43]. In distant-site TADE suppression, the version of the drive that we model in this study, female fertility gene disruption is achieved by locating the drive itself inside the female fertility gene (not via gRNA targeting). This means that the haplolethal target gene that the drive cleaves is at a distant site to the drive. Because drive homozygous females are sterile, strong population suppression is achieved with this drive. However, it still only has a nonzero introduction threshold if there are fitness costs or cleavage in the embryo form due to maternal Cas9/gRNA deposition. Our ideal TADE suppression drive has no fitness costs, 100% germline cleavage, and no embryo cleavage. All the possible genotypes in our model are listed in Figure 1C.

**CifAB** utilizes the toxin-antidote mechanism of two genes in prophage WO of *Wolbachia*: *CifA* and *CifB*. It thus has the same cytoplasmic incompatibility mechanism as *Wolbachia*, but on the genetic level rather than with a bacterial infection [38]. A male with the *CifB* gene can cause cytoplasmic incompatibility (the toxin), while a female with the *CifA* gene (sometimes also requiring the *CifB* gene) produces an antidote [44–46]. The absence of the antidote in the presence of the toxin prevents offspring from being produced (Figure 1D). For this drive, we assume that *CifA* and *CifB* are both in the drive allele. We assume that all crosses produce normal offspring, except wild-type females crossed with any drive carrier male, which produces no offspring. This “CifAB drive” is a highly confined system with an introduction threshold of 37% [38, 41] in the ideal form that we model (100% cytoplasmic incompatibility and no fitness costs).

***Wolbachia* bacteria** can spread through a population in a similar manner to a gene drive [32]. They are vertically transmitted by infected females to all their progeny, whereas infected males do not transmit the bacteria. However, cytoplasmic incompatibility results in no offspring if wild-type females mate with *Wolbachia*-infected males. *Wolbachia* exhibits an introduction threshold whenever infection imposes a fitness cost. In this study, rather than an ideal system, we use a fitness of 0.75 (affecting female fertility and male mating success) for *Wolbachia*-infected individuals based on field observations and experiments [47]. This gives *Wolbachia* an introduction threshold of approximately 31%.

### Reaction-diffusion system

We use a reaction-diffusion system to simulate population dynamics:

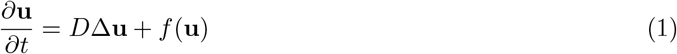

here **u** is the vector of population sizes for each genotype, *D* is the dispersal coefficient (the average dispersal, set to 1 by default), and *f* (**u**) is the reaction term. Assuming radial symmetry, the Laplacian 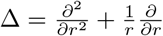. The reaction term is as previously reported [48]:

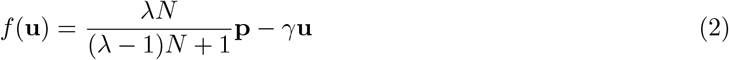

where *N* = ∑*u*_*i*_ is the total population size, **p** is a vector of probabilities, and its *j*-th entry is the probability of getting an offspring of genotype *j* given the current population composition including type of gene drive or *Wolbachia* system (see supplemental section eqs. (8), (9), (10), (11) and (12) for details). *λ* is the intrinsic growth rate (by definition one lower than the relative “low-density growth rate” used in some other models), and *γ* is the death rate (set to 1 by default, so 1 is the generation time at equilibrium). Model parameters are listed in Table 1.

**Table 1.** Parameters used in the reaction-diffusion system.

| Variable | Name | Value |
| --- | --- | --- |
| $D$ | Dispersal coefficient | 1 |
| $\lambda$ | Intrinsic growth rate | 9 |
| $\gamma$ | Death rate | 1 |

### Release pattern and spread efficiency

In general, a release pattern can be any function supported on a two-dimensional area. However, in real-world scenarios, executing a precisely variable release pattern is often impractical due to complexities in topography and transportation logistics. Furthermore, optimizing over a two-dimensional variable release shape is computationally expensive. With these two considerations, we simplify the problem to finding the optimal release pattern among all patterns that are centered at an origin and are radially symmetric (Figure 2). This will contain the actual optimal release pattern while minimizing the computational intensity of the problem.

**Figure 2.**
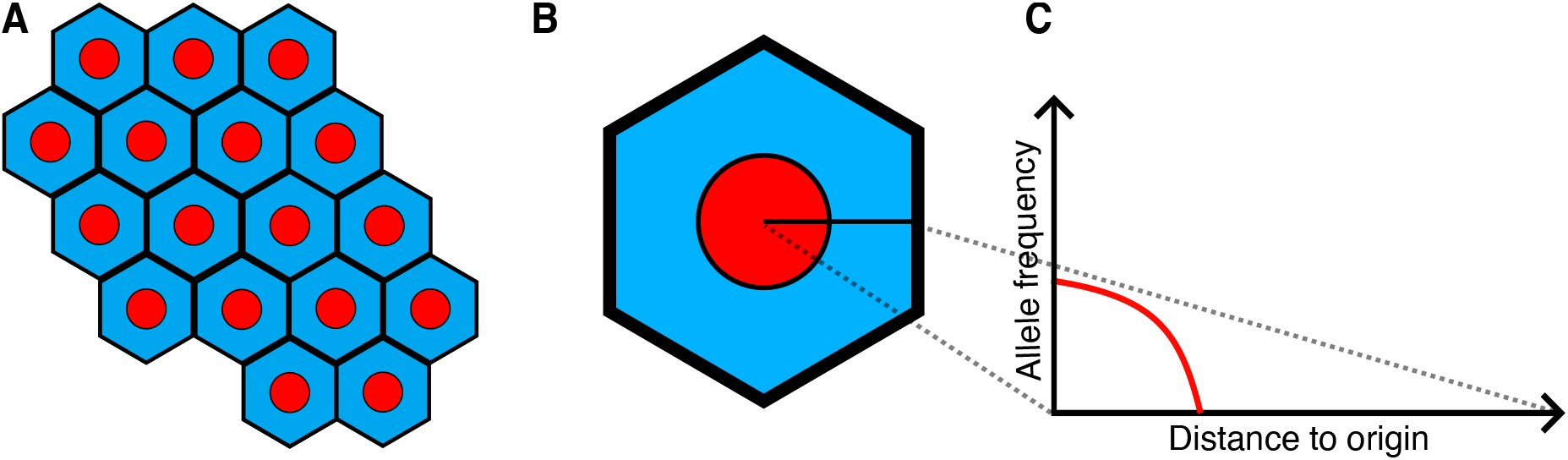
Schematic of release strategy. **A** A larger region of wild-type individuals (blue) to be controlled by multiple gene drive releases (red). Each release is responsible for spreading the gene drive into one hexagon (with possible contribution from the surrounding drive releases). **B** A view of one gene drive release site. The gene drive release is centered at an origin and is radially symmetric. **C** The gene drive release frequency as a function of radius in the single release zone.

**Figure 3.**
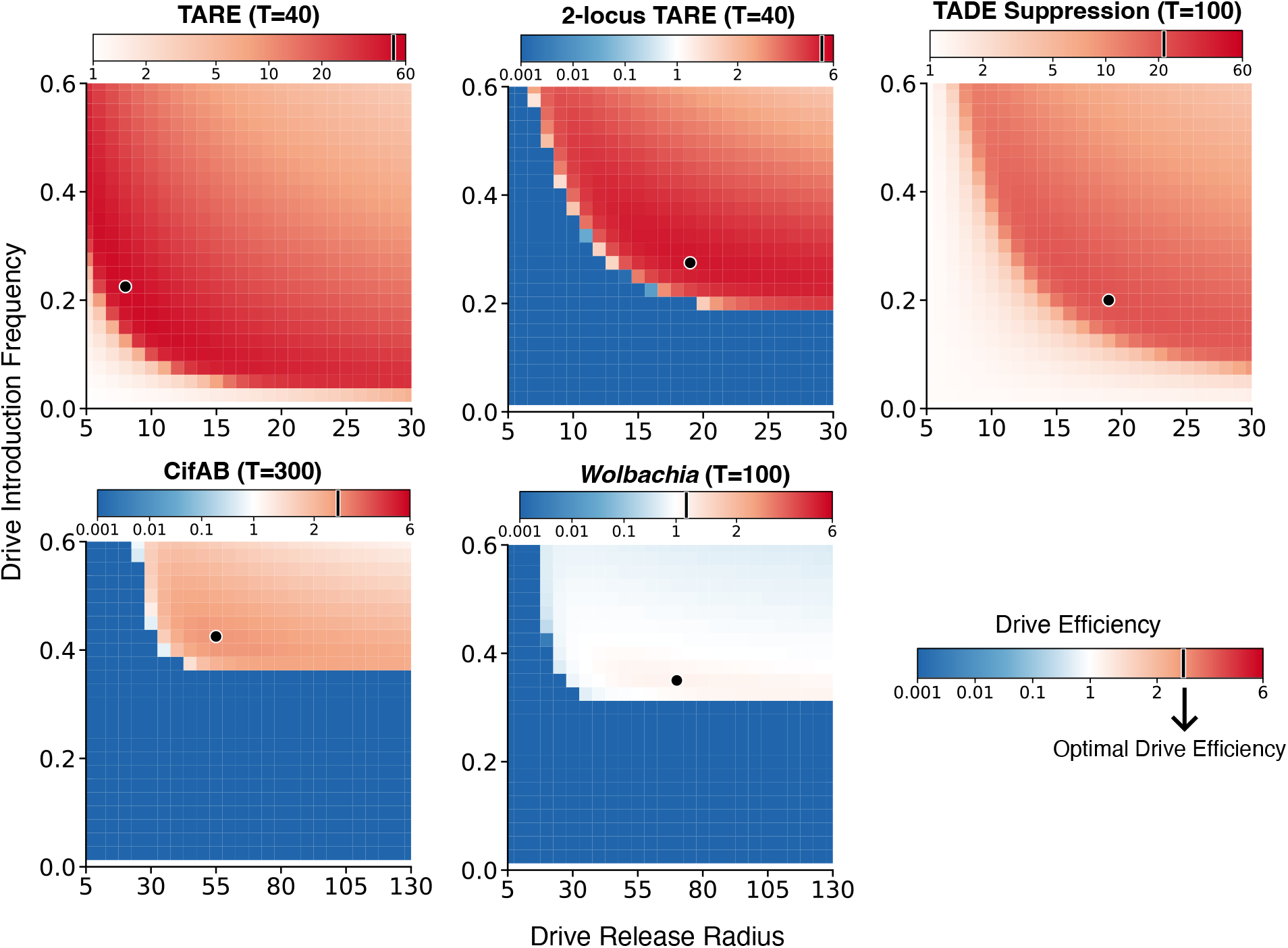
Spread efficiency for circular releases. Simulation times are chosen as indicated for different systems. Each panel shows spread efficiency for a circle release as a function of release frequency and release radius. Black points indicate the most efficient release within each heatmap. Note the different scales for efficiency and release radius for different systems.

Under the radial symmetry assumption, the problem is 1-dimensional. We drop drive individuals at time *t*_0_ = 0, which serves as the initial condition:

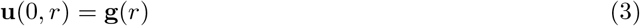

We only release drive individuals of a single genotype, say genotype *i*. Therefore, all but two entries in **g** are zero (the population size of wild-type individuals starts at its equilibrium of 1). We refer to the *i*-th entry of **g** as the release pattern:

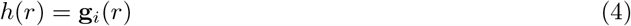

We will optimize this function.

Now we introduce the objective function in our optimization, the spread efficiency of a gene drive. For modification drives (TARE, 2-locus TARE, CifAB and *Wolbachia*), we define spread efficiency as:

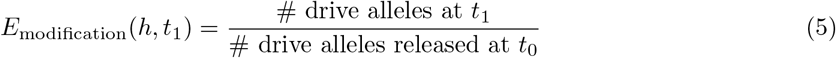

For suppression drives (TADE suppression), spread efficiency is:

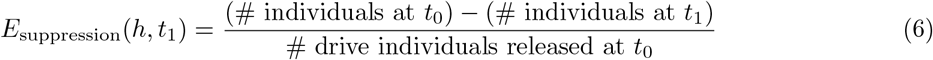

In other words, the spread efficiency of a modification drive is the number of drive alleles we can obtain on average by releasing one drive allele, and the spread efficiency of a suppression drive is the number of individuals we can eliminate on average by releasing one drive individual.

### Numerical computation

We use the explicit finite difference method for numerical simulation of the reaction-diffusion equation. The radius of the whole arena *R* is set large enough so that the wave never reaches the boundary of the whole arena (*R* = 400 for all sections except the long term release pattern section, which uses *R* = 10000). The maximum radius of the release area is kept at *L* = 200. A spatial-temporal mesh of Δ*x* = 0.5 and Δ*t* = 0.05 is used throughout. While a finer spatial mesh would improve resolution, the Von Neumann stability criterion for heat equations requires the time step to scale quadratically with the spatial step (Δ*t* ∼ Δ*x*^2^) to avoid numerical instability. This means that the computational cost scales cubically with the inverse of the spatial resolution (*O*(Δ*x*^−3^)). We arrive at Δ*x* = 0.5 because further reduction in Δ*x* would result in a prohibitive increase in runtime for the iterative optimization process. We find that this resolution produces sufficiently smooth curves for our variable shape release patterns.

#### Optimal constant-density circle releases

We first restrict the optimization to circle releases that have the same release density throughout the circle. A circle release is parameterized by the release radius *r*_0_ and release frequency *f*_0_.

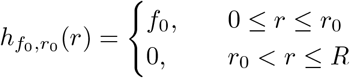

We investigate this for two reasons. (i) In some real-world scenarios, it may be difficult to release the gene drive in a complicated shape, while a constant-density release in a single area is relatively simple. (ii) The drive spread efficiency of an optimal circle release serves as our baseline for the drive spread efficiency of the optimal variable release discussed in the next section. We denote the set of all circle release patterns as ℋ_circle_, and for a set of times 𝒯, we find the corresponding optimal circle release pattern:

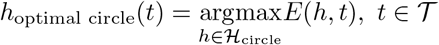

#### Optimal variable releases

We next extend the optimization to variable release curves. Under our spatial mesh, a variable release pattern is any piecewise constant step function on the interval [0, *L*].

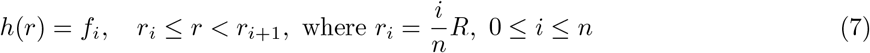

We denote the set of variable release patterns as ℋ_variable_, and for a set of times 𝒯, we find the corresponding optimal variable release pattern:

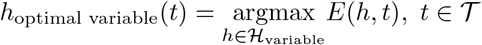

### Data generation and software

Simulations and optimizations were performed in MATLAB R2025b. All code used in this study is available at https://github.com/Zii-yyE/gene-drive-optimal-spatial-release.

## Results

### Drive efficiency of circular releases

Before comparing optimal release strategies, we first asked how much spread efficiency can vary between circle releases. To do so, we calculated the spread efficiency (see methods) of circle releases with varying release radii and introduction frequencies at an intermediate timeframe for each drive system where the optimal release would not be an “everywhere” release (see below). The resulting heatmaps reveal that spread efficiency is highly sensitive to the release pattern. Even under this restricted class of releases, different choices of radius and frequency can lead to markedly different outcomes, ranging from highly efficient spread to poor performance, and in some systems spanning several orders of magnitude. Releases below the introduction thresholds for 2-locus TARE, CifAB, and *Wolbachia* always failed, of course, and releases close to the failure threshold also had poor performance. However, if releases were too wide or too high, efficiency also suffered. The highest efficiency would usually found in a band moderately above release requirements in both radius and release frequency. We can thus conclude that optimization of the drive release pattern can substniually improve the overall efficiency of a drive release.

### Optimal circle release pattern

We performed optimization on circle release patterns for TARE, 2-locus TARE, TADE suppression, CifAB, and *Wolbachia* bacteria (Figure 4, see also Figures S1 to S5). In most cases, we used ideal drive forms, but for *Wolbachia*, we used a fitness cost of 0.25 to match real-world releases (see methods). The optimal pattern for modification drives is one that results in the most alleles at the final timepoint for each drive allele released. For the suppression drive, the optimal pattern is the one that results in greatest population reduction for each drive individual released. The optimal circle release pattern changes based on the final timepoint. With a short timeframe, the optimal circle pattern will cover the whole area allowed for release (note that this doesn’t represent the entire arena in this model, but the optimal release would indeed cover the entire region). This is because the drive does not have time to form a wave and advance and spread to new areas. It can only somewhat increase in frequency in the release area. As time increases, the radius of the optimal circle pattern suddenly shrinks to no longer occupy the whole release area, while the release frequency within the release area somewhat increases. As the time further increases, the radius of the release gradually decreases and the introduction frequency becomes stable, usually but not always increasing asymptotically.

**Figure 4.**
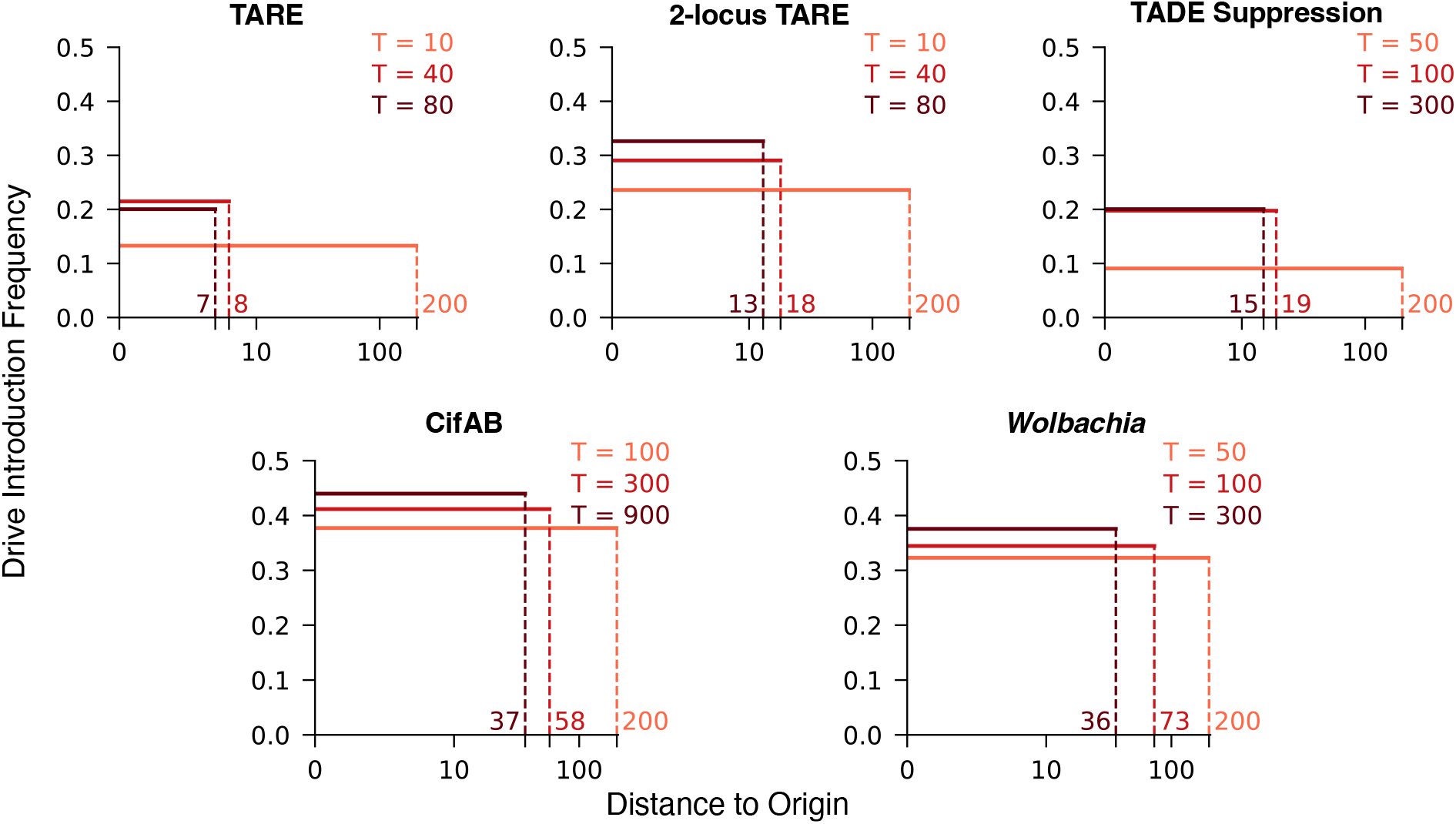
Optimal circle release pattern. Graphs show the optimal release radius and frequency for three total simulation times (in generations) for each drive (note that different drives have different simulation times displayed). Units of distance correspond to the average dispersal in one generation. The maximum possible release radius is 200.

As expected, the more powerful drives required lower release sizes (Figures S6 and 7) and release radii (Figure 4, Figures S1 to S5), also transitioning more quickly from “everywhere” releases to releases in a restricted region. TARE did this most quickly, followed by 2-locus TARE and TADE suppression. Note that despite the lower introduction threshold of TADE suppression compared to 2-locus TARE, it is still a slower drive due to its suppression effect [41]. These were followed by *Wolbachia*, which was substantially slowed by its fitness cost, and finally by CifAB drive, which has a high threshold and slow wave of advance.

### Optimal variable release pattern

We next fully optimized the release patterns for each drive, allowing for a variable release shape across the entire release area (Figure 5, see also Figures S1 to S5). The pattern also changes with time, going through several stages. For the shortest timeframes, the optimal pattern is an “everywhere release”, similar to the circle releases. However, the release density tends to be slightly larger near a radius of 200, which is a minor artifact of this maximum radius. This higher release near the boundary between drive and wild-type is also seen in some other release patterns for systems with higher introduction thresholds. It allows for higher drive density where dispersion may otherwise reduce drive density due to proximity of nearby wild-type, thus allowing the drive to remain above its introduction threshold and continue to spread. For relatively longer timeframes, the optimal pattern smoothly but quickly transitions to a “multiple-ring release” (with a central release zone included in some cases). Unlike the “everywhere” release, the multiple-ring release covers only some of the territory using several rings of drive release and leaving wild-type rings between them. This occurs when the drive is able to slightly spread out due to dispersal in the simulation timeframe, covering all the space between the release rings. For even longer timeframes, the optimal variable release eventually transitions to a single ring (which will cover the middle area within the simulation timeframe) and ultimately to a single central release. With these longer times, the gene drive has sufficient time to form a full wave of advance and spread a substantial amount, so it is no longer necessary to release it across the whole region. A single release covers less territory, but multiple releases could be used to efficiently cover the whole arena if needed. The drive wave advance now significantly increases the amount of drive alleles present in not just the release area, but well outside of it.

**Figure 5.**
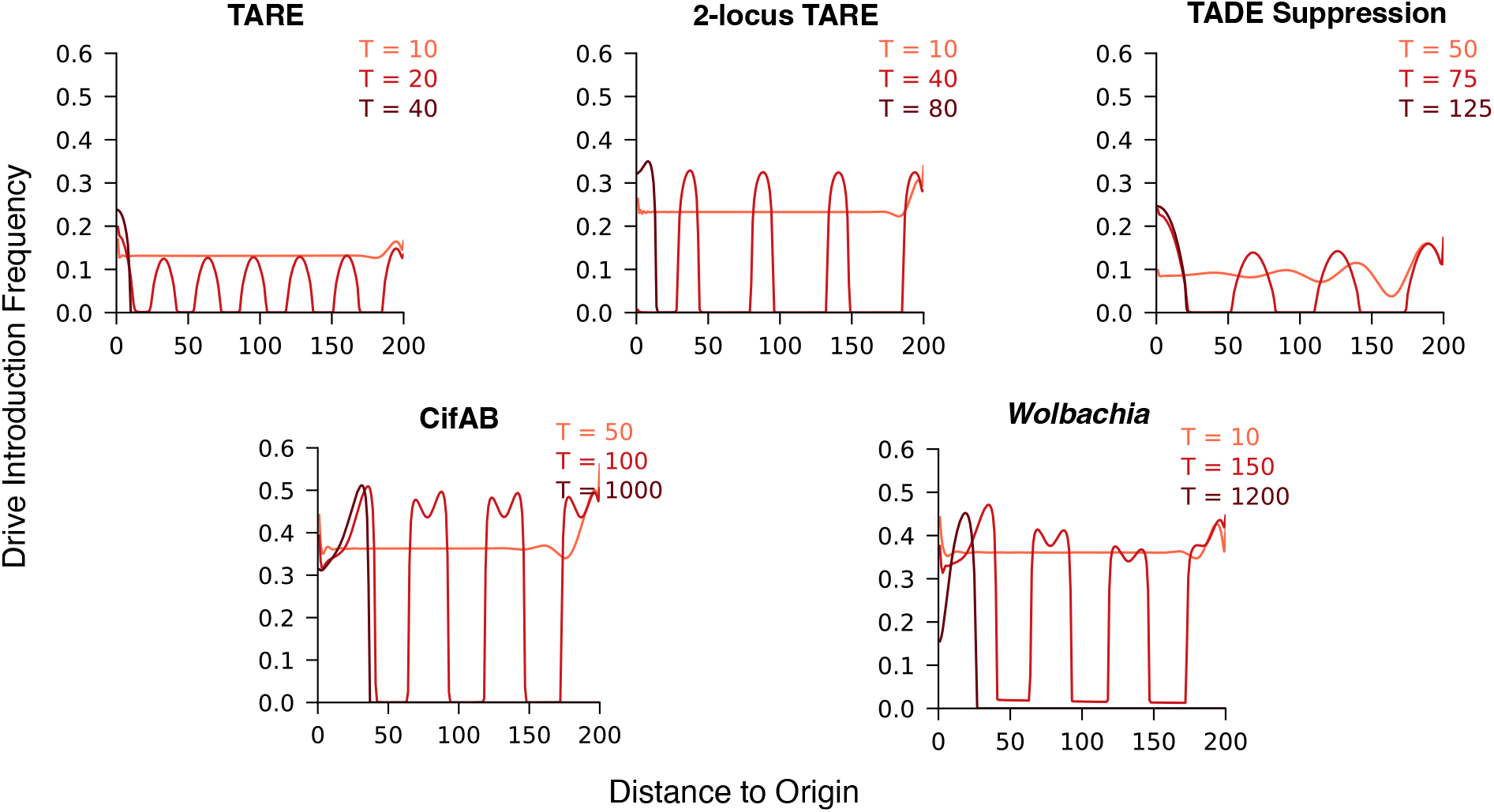
Optimal variable release pattern. Graphs show the optimal release radius and frequency for three total simulation times (in generations) for each drive (note that different drives have different simulation times displayed to show the different release pattern forms). Units of distance correspond to the average dispersal in one generation. The maximum possible release radius is 200.

### Long term optimal release pattern

As the timeframe of the simulation increases to an arbitrarily large level, the drive will expand outward, forming a circle. In a small circle, surrounding wild-type will cause the drive to be at a geometrical disadvantage due to its frequency-dependent performance [21], slowing its advance. As the circle expands, though, this effect will be reduced. Eventually at the edge of the circle, it will advance nearly as quickly as a flat wave of drive advancing into wild-type, asymptotically reaching this peak speed. When this phase of drive advance dominates the timeframe, then further increases in the timeframe should not significantly affect the optimal release pattern. This is consistent with our results thus far, where we observed that the optimal release pattern does not substantially change for longer times for higher threshold drives. We thus sought to assess this long-term optimal release pattern by optimizing our drive release for a very long time window (*T* = 10000). This was performed both for circular constant-frequency releases and variable pattern releases (Figure 6). In all these releases, the frequency is about 5-20% above the threshold over most of the release area. The lowest threshold drives have the smallest total release size, despite 2-locus TARE having a greater wave advance speed than TADE suppression.

**Figure 6.**
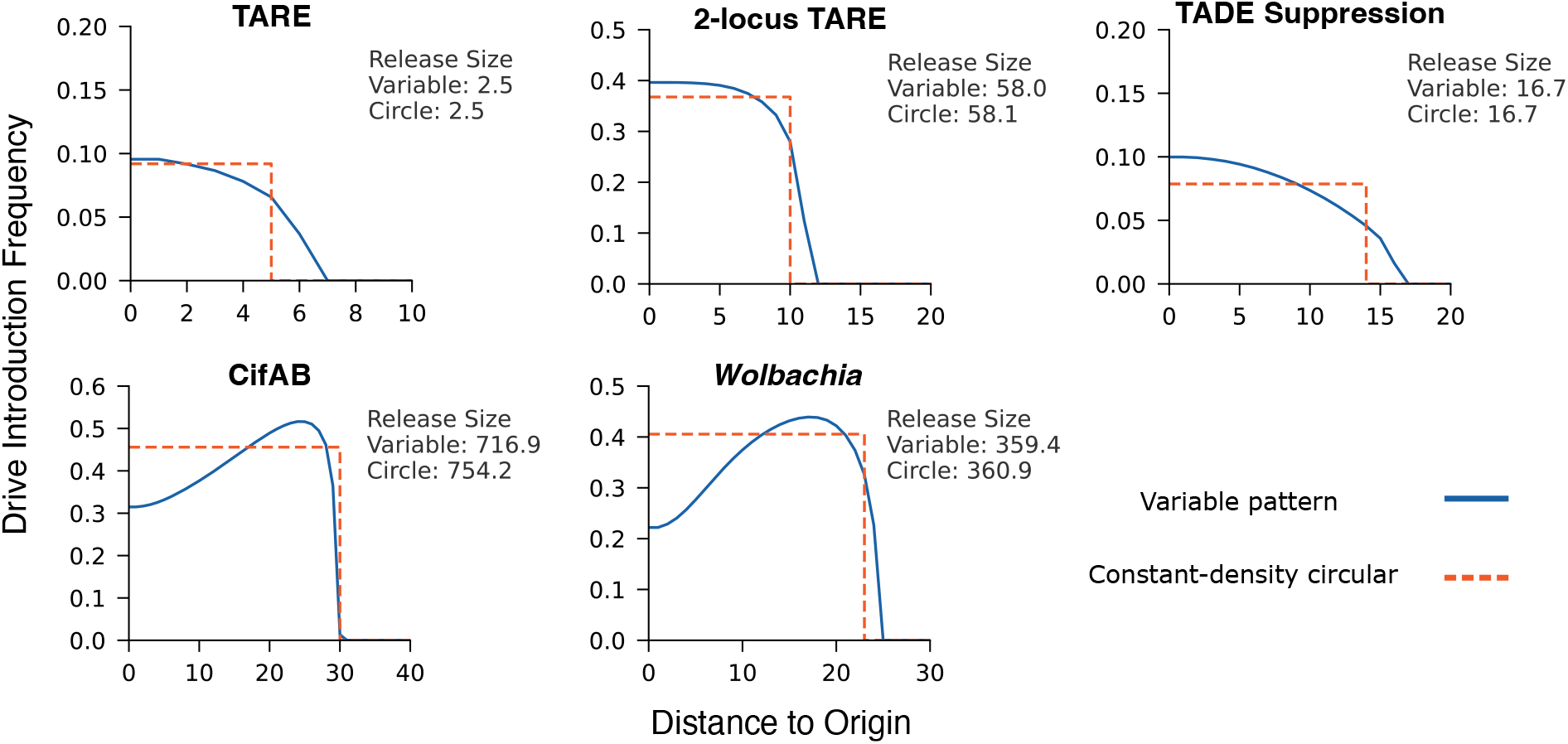
Long term optimal release pattern. Graphs show the optimal release radius and frequency for both circular constant-density and variable release patterns for each drive. All drives have the same simulation time *T* = 10000 generations. Different drives have different axis scales. Units of distance correspond to the average dispersal in one generation.

We note that the constant-density circular releases and variable releases are highly similar in shape and usually have a less than 1% difference in release size (CifAB with the highest threshold has a less than 5% difference). For the lower threshold systems, the variable release pattern has a higher concentration in the center to allow for a strong core of drive establishment, followed by outward expansion. For the higher threshold systems (CifAB and *Wolbachia*), the optimal shape has lower density near the very center, which will be covered by outward migration for further toward the edge of the release circle. The outer region of the release circle has a release frequency well above the introduction threshold, allowing it to withstand dispersal from wild-type individuals surrounding the circle, thus helping to overcome the geometrical disadvantage from a central release.

### Optimal spread efficiency

We next examined the spread efficiency achieved by optimal release patterns (Figure 7). The release size, if not overall efficiency, in these strongly correlated with the general class of release pattern (everywhere, multiple ring, central release). For short timeframes, the highest efficiency is achieved by releasing a large number of drive individuals, which corresponds to the “everywhere release”. As the timeframe declines, the number of released individuals drops rapidly in “ring release” patterns, with the rings becoming more spread out. This particular phase is not present for constant-density circular releases. After a critical point in time, the release size drops to a low level and afterward only modestly declines (Figure S6), representing a single central release.

**Figure 7.**
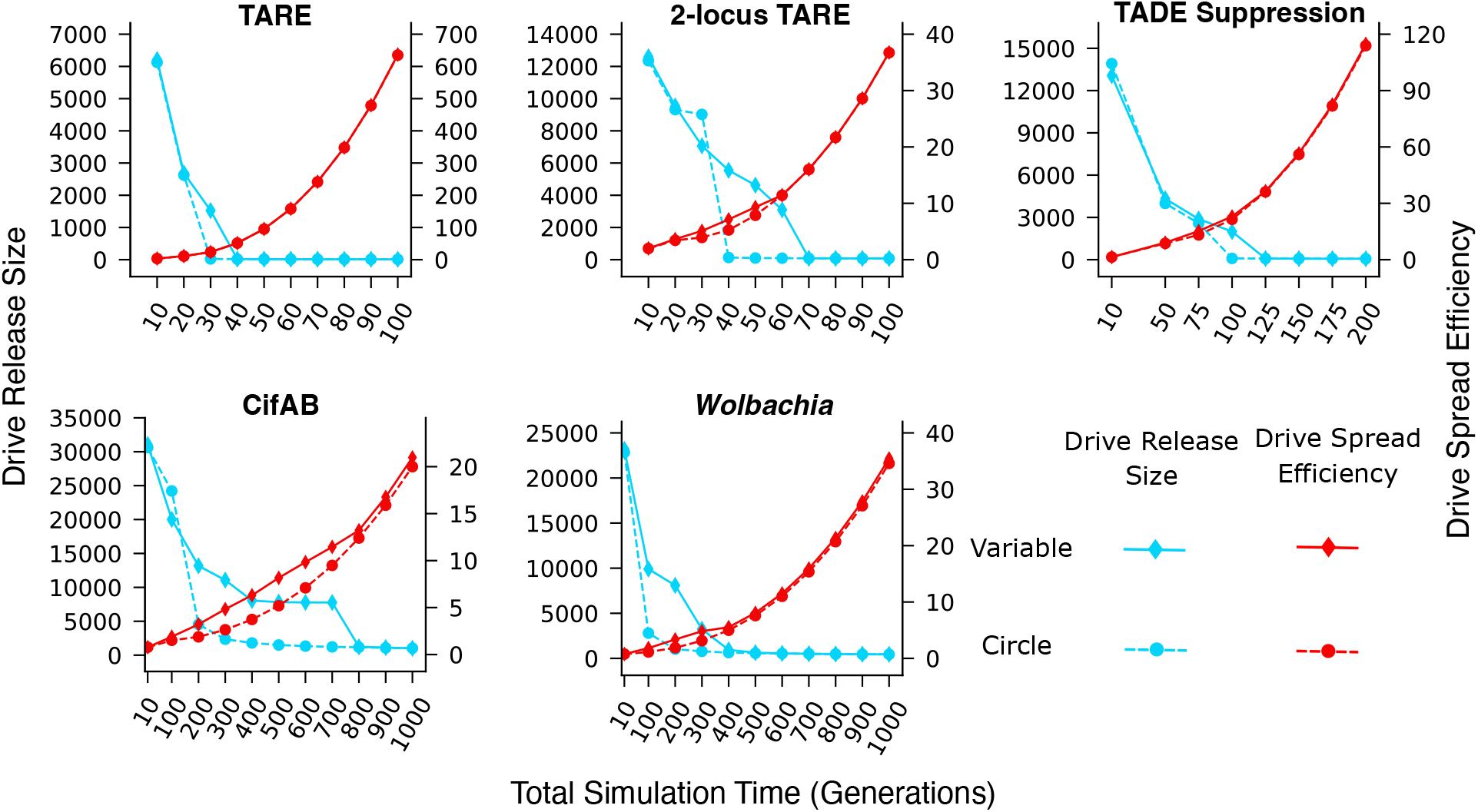
Drive spread efficiency and release size. For constant-density circle and variable optimal drive release patterns, we show the total release size on the left axis, which is the fraction of wild-type individual density in one unit area, defined by the average dispersal distance. On the right axis, we display drive spread efficiency. Figure S6 more clearly shows release sizes at larger timescales for each drive.

The type of drive plays a significant role in overall efficiency. In general, the overall efficiency is correlated with the introduction threshold, though TADE suppression is the exception again, with its slower wave advance reducing efficiency compared to 2-locus TARE, despite its lower release level. The faster drives also more quickly transition from an everywhere release to a single point release. Interestingly, with optimized releases, the efficiency does not always strongly correlate with release pattern (constant-density vs. variable, everywhere vs. ring vs. single central). In TARE drive, for example, the shape of the efficiency curve does not significantly deviate between low and high timescales, and the variable release pattern is only marginally more efficient than the constant-density circle release. For the other drives, variable vs. circle efficiency is also similar at higher timescales (single central release) and very short timescales (everywhere release). In between, the ring release in the variable pattern can provide a substantial efficiency improvement (in particular CifAB, and only marginally for TADE suppression) compared to a constant-density circular release. For these drives, the variable release efficiency tends to linearly increase, before eventually following a square law at higher efficiency (in the long term, the drive circle expands its radius at a near-constant rate). This is in contrast to the smoother efficiency curves for the constant-density circular release.

### Drive efficiency compared to panmictic models

In some cases, it may be possible to do a widespread gene drive release, or in some species, populations may be better represented as panmictic. In these cases, optimized gene drive release is simplified, requiring only a single parameter: the introduction frequency (which will be above its threshold, with the exact amount depending on the timescale). In panmictic models, there is also a maximum gene drive frequency, rather than the potential to continue expanding in spatial models of large arena. This provides a maximum possible efficiency for threshold-based drives, which must be released above the threshold to succeed. On the other hand, spatial models create a more difficult situation for the gene drive at first, with dispersal from wild-type outside the release areas. We were thus interested in comparing drive spread efficiency between optimal spatial and panmictic releases.

To assess this, we ran simulations to optimize drive release in the panmictic model and calculated drive spread efficiency in the same way as in our spatial models (Figure S7). We then plotted the ratio of drive spread efficiency from our variable release in the spatial model to the same efficiency in the panmictic model (Figure 8). At first, the pattern shows lower efficiency in the spatial models, especially for TADE suppression, which is particularly vulnerable to frequency-dependent effects. This is the region of the “everywhere” release in the spatial model, but the maximum release area is still smaller than the total arena size. Thus, wild-type dispersal still reduces efficiency compared to the panmictic model, albeit only a small amount. However, with an expanded timeframe, the spatial release changes to other patterns, and drive spread to new areas allows efficiency to be substantially higher in spatial populations than for panmictic populations.

**Figure 8.**
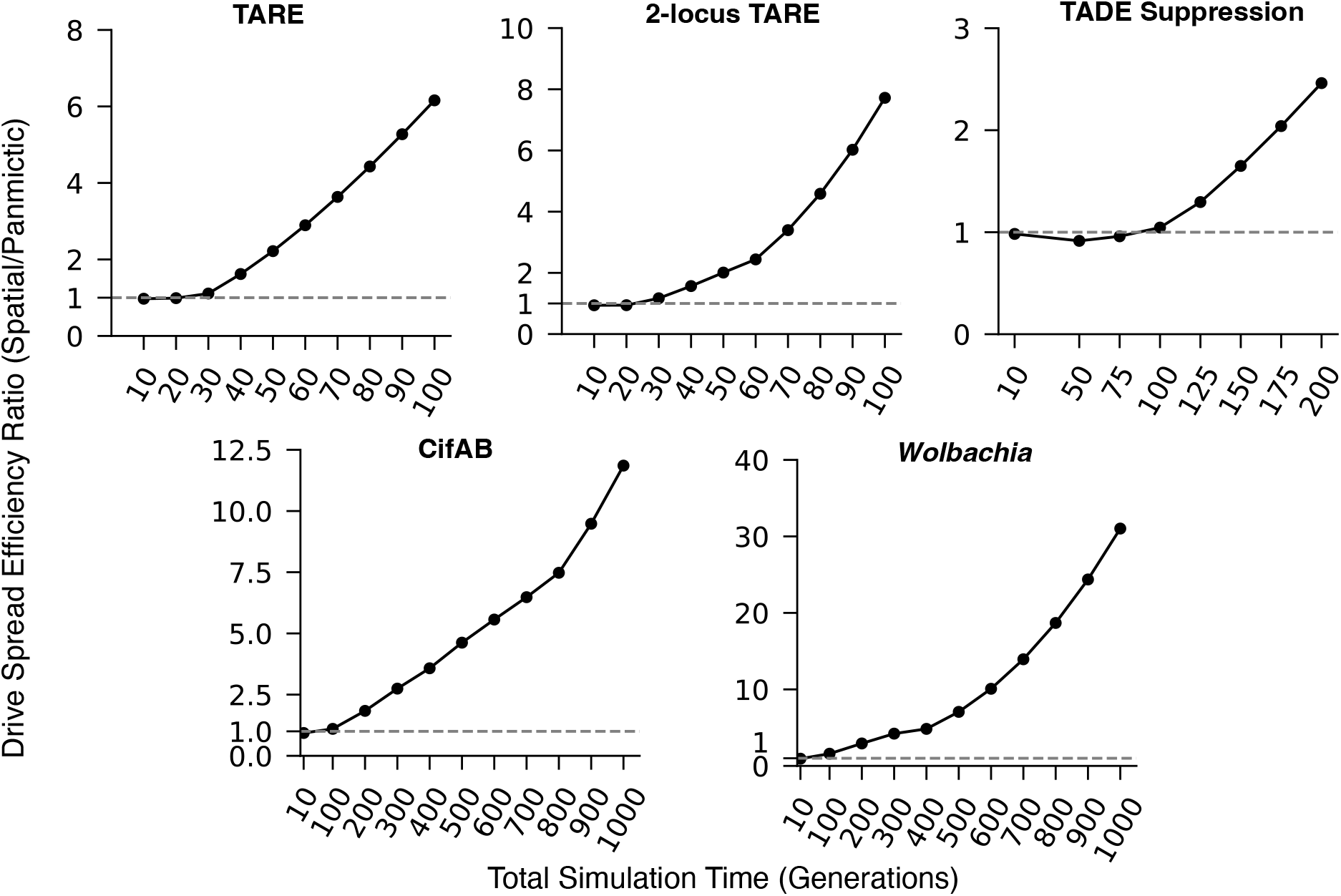
Drive efficiency comparison between spatial and panmictic models. The plots show the efficiency ratio for each system as a function of the simulation timeframe. This is the drive spread efficiency for the optimal variable release pattern in the spatial model divided by the drive spread efficiency for the optimal release level in a panmictic model.

## Discussion

In this study, we optimized release strategies for four different gene drives—TARE, 2-locus TARE, TADE suppression, and CifAB, plus *Wolbachia*, using a reaction-diffusion model. Our goal was to identify release patterns that maximize drive spread efficiency over a given timeframe. Results reveal that the optimal release strategy depends critically on both the type of the gene drive and the intended operational timeframe. A key finding is that optimal release strategies follow several distinct patterns: an “everywhere” release for short timeframes, a “multiple-ring” release for intermediate times, and a central release for longer timeframes. This transition from broad coverage to a focused core reflects a strategic logic that leverages the drive’s own dispersal capabilities to maximize the effect per released drive individual. The “multiple-ring” pattern is a particularly non-intuitive and powerful strategy, effectively seeding a large area and relying on dispersal to fill the intervening gaps, thereby achieving control with a smaller investment of engineered organisms.

While we did not compare optimal release strategies to a variety of non-optimal patterns, previous work has shown that substantial efficiency advantages are accrued [38], making these optimal releases patterns valuable. Compared our optimal circular and variable pattern releases, we see that the degree to which different drives benefit from the variable release pattern. For a highly efficient drive like TARE, the performance gain from a variable release over a simple optimized constant-density circle release is modest. While such an optimized release can still be better than alternatives, there will usually be no need for more advanced strategies for such a drive. In contrast, for confined drives with higher introduction thresholds, such as 2-locus TARE and CifAB, an optimized variable release can be significantly more efficient than the best circle release unless the timeframe for the intervention is very long. This highlights a critical principle: for drives that are inherently difficult to establish and spread, a well-planned spatial release strategy is beneficial and may be essential for success.

In analyzing the release pattern, we noted that for the higher threshold CifAB drive and *Wolbachia*, the optimal release frequency is lower near the origin and higher toward the edge of the release area. This represents an efficient strategy that concentrates release efforts at the propagating wave front, which will suffer from an initial influx of wild-type individuals during initial dispersal. However, we did not observe this pattern for the central releases of lower threshold drives. These systems are still frequency dependent, but their lower threshold means that the edges are in far less danger of being overwhelmed by influx of wild-type alleles. This effect still slows the drives, but outward migration is enough to stabilize the initial condition without needing a higher frequency of drives on the edge of the release area. Interestingly, we did observe this release pattern for central releases of 2-locus TARE for intermediate times, but it disappeared when the timeframe became larger. This may be because after a sufficiently long time, the optimal release does converge to a critical bubble. This is why the zero-threshold drives (with a zero-size critical bubble) continue to slowly reduce release size after an extended period of time, while the higher threshold drives asymptotically converge on the critical bubble.

This work potentially fills a useful niche in analysis of frequency-dependent gene drives and other genetic population engineering systems. While previous studies have investigated strategies for releasing gene drives, they have mostly focused on unconfined systems such as homing drives. Those that evaluated frequency-dependent drives usually did not investigate continuous space models (or models producing analogous results), and those that did focused mainly on the critical release, which is different from the optimal release we seek in a real-world scenario, at least for most meaningful timescales. Even panmictic studies tend to focus on thresholds rather than optimized releases for a specific timeframe. Our study also investigates some promising new types of drives, which have different spatial properties than systems assessed in older studies [40, 41]. Moreover, previous studies have often been limited to comparing a few simple, pre-defined release patterns [16, 38]. Our study systematically optimized the release pattern over a continuous, radially symmetric space, providing a more rigorous answer to the question of how to best release a drive. Our findings support the notion that spatial dynamics are of paramount importance to the population dynamics of confined gene drives.

While a potential starting point for planning a gene drive release, our study necessarily makes several simplifying assumptions. We assumed radial symmetry, which neglects the environmental heterogeneity of real-world landscapes, which can influence density and have a large effect on spread [21, 32, 49]. Our model is also deterministic and assumes ideal drive characteristics, such as 100% drive efficiency, no fitness costs, and the same dispersal coefficient everywhere in the landscape. Stochastic events in particular may be highly important when predicting the outcome of a suppression drive [42, 50, 51]. These considerations can influence drive performance and should be considered [38, 40]. Individual-based, stochastic models may be crucial for validating their applicability to real-world scenarios, where chance events can play a significant role at the low-density wave front. Extending this optimization framework to spatial models with heterogeneous landscapes would be a valuable next step. Furthermore, future work could explore alternative optimization objectives, such as minimizing the time to modification/suppression for a fixed release budget or vice versa. Logistical costs could also be incorporated if such data were available. Even in a simple situation, with a maximum release level in a certain timeframe, but with possible ongoing releases, the optimization problem would become considerably more complicated, especially if the available release capacity was insufficient to form at least one or more optimized release patterns in a single release event. Indeed, it might be that a release over a few generations is more optimal for high threshold drives because of less reproduction loss at high densities. For a specific real world application, use of region-, species-, and drive-specific models should be evaluated together with these considerations for determining if a gene drive release is a suitable strategy and how such a drive should be deployed.

In conclusion, this study demonstrates that the spatial arrangement of a gene drive release is a critical and optimizable parameter. Suitable strategies, such as the multiple-ring release, can increase the efficiency of a drive release over simpler circular releases, particularly for highly confined gene drives. This work underscores the necessity for careful strategic planning for field implementations of gene drive technology, where an optimized release pattern could mean the difference between success and failure.

## Acknowledgments

This study was supported by the Center for Life Sciences and the National Natural Science Foundation of China (grants 32270672 and W2432018).

## Supplemental Information

### Supplementary Figures

**Figure S1.**
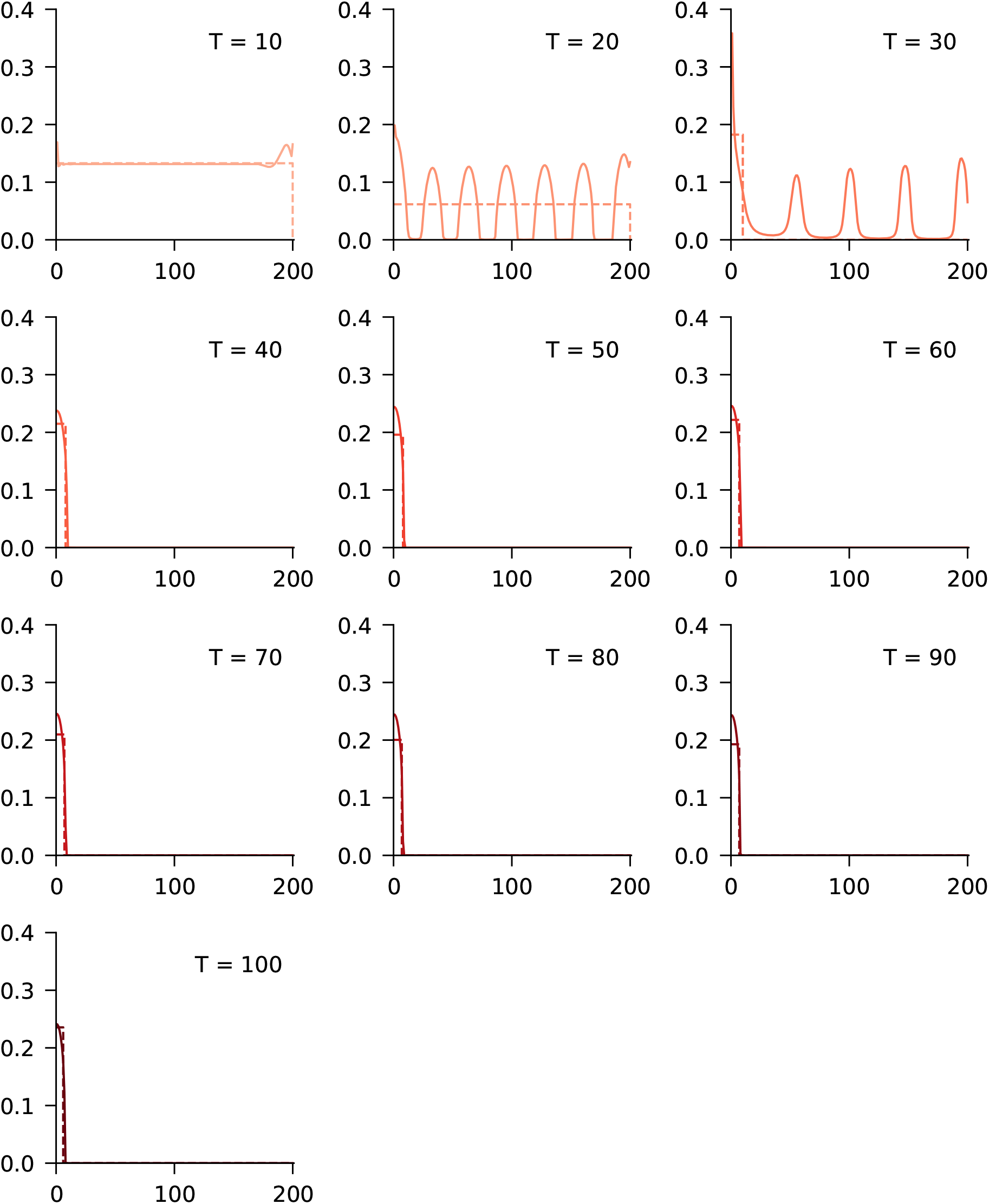
Optimal release patterns for TARE. Solid lines are optimal variable release patterns. Dashed lines are optimal circle release patterns. T shows the number of generations for the optimization timeframe.

**Figure S2.**
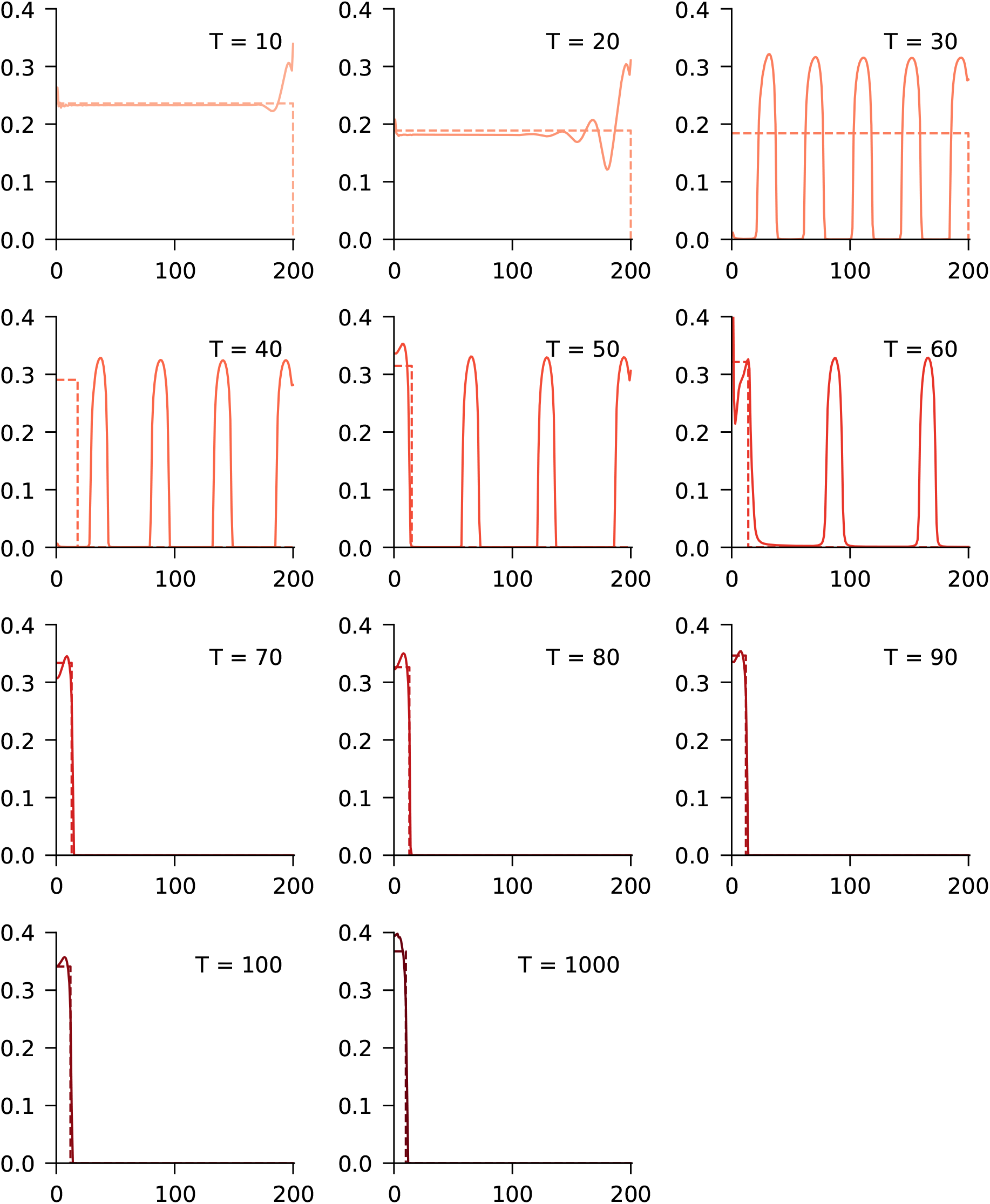
Optimal release patterns for 2-locus TARE. Solid lines are optimal variable release patterns. Dashed lines are optimal constant-density circle release patterns. T shows the number of generations for the optimization timeframe.

**Figure S3.**
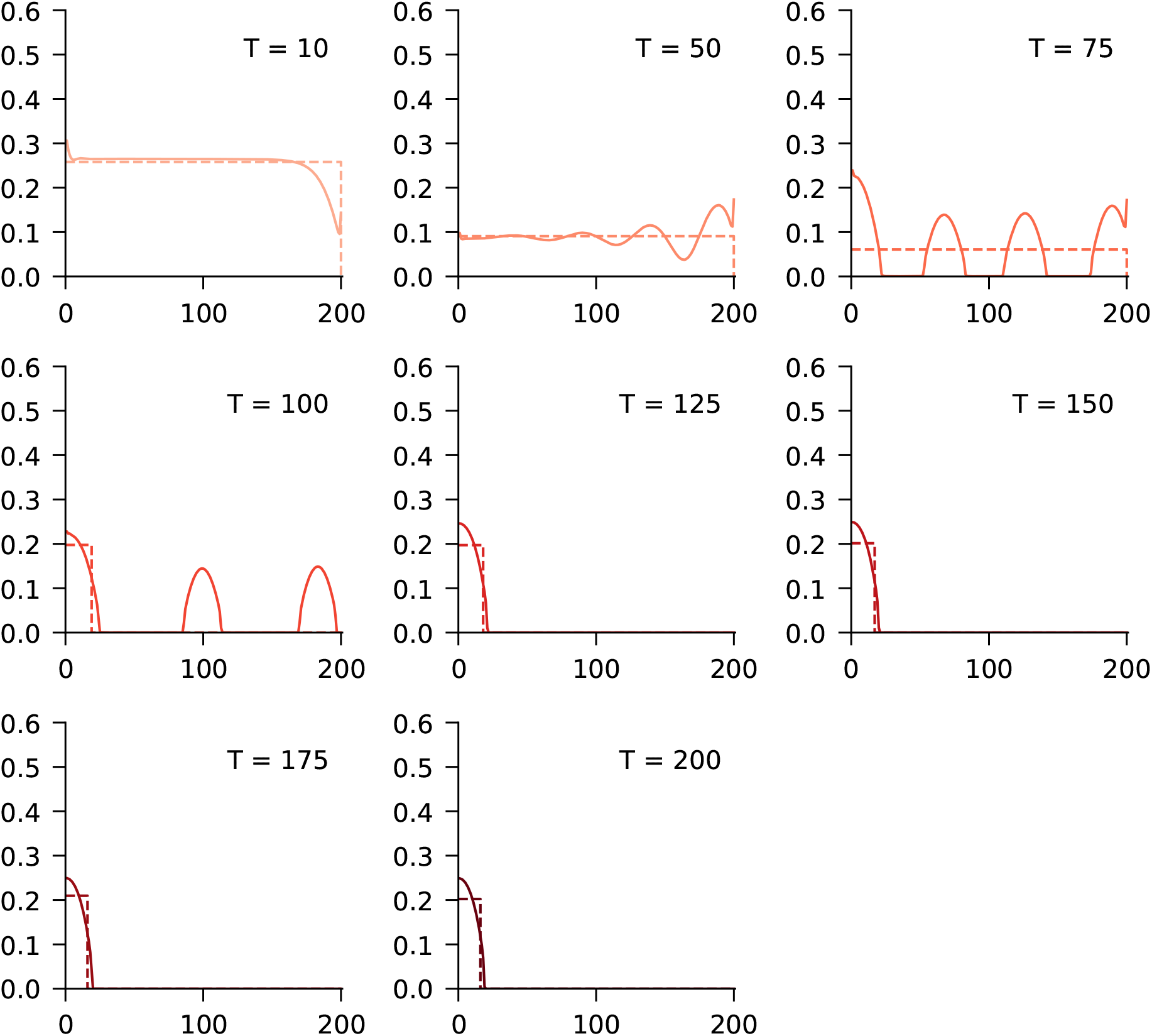
Optimal release patterns for TADE suppression. Solid lines are optimal variable release patterns. Dashed lines are optimal constant-density circle release patterns. T shows the number of generations for the optimization timeframe.

**Figure S4.**
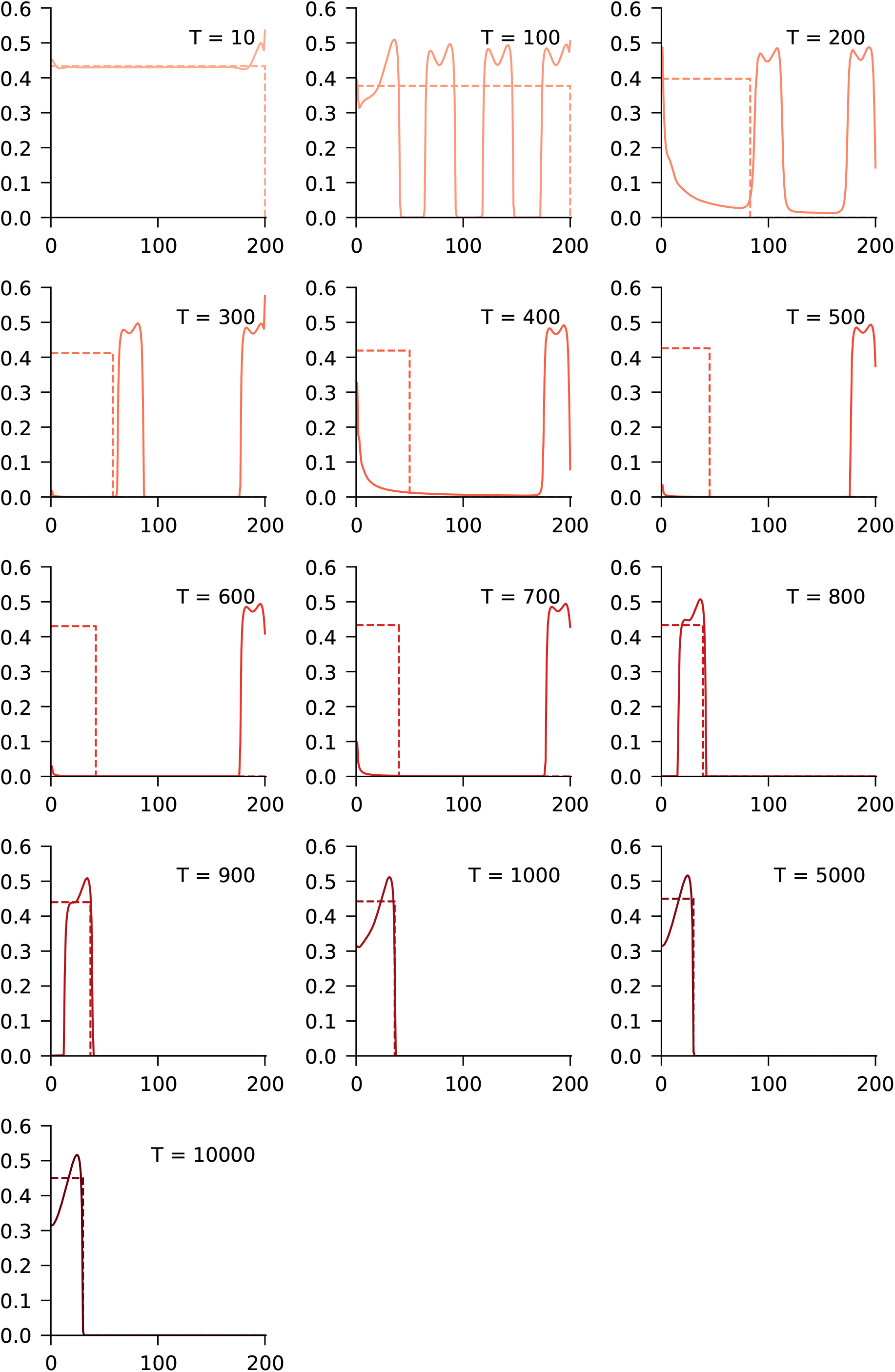
Optimal release patterns for CifAB. Solid lines are optimal variable release patterns. Dashed lines are optimal constant-density circle release patterns. T shows the number of generations for the optimization timeframe.

**Figure S5.**
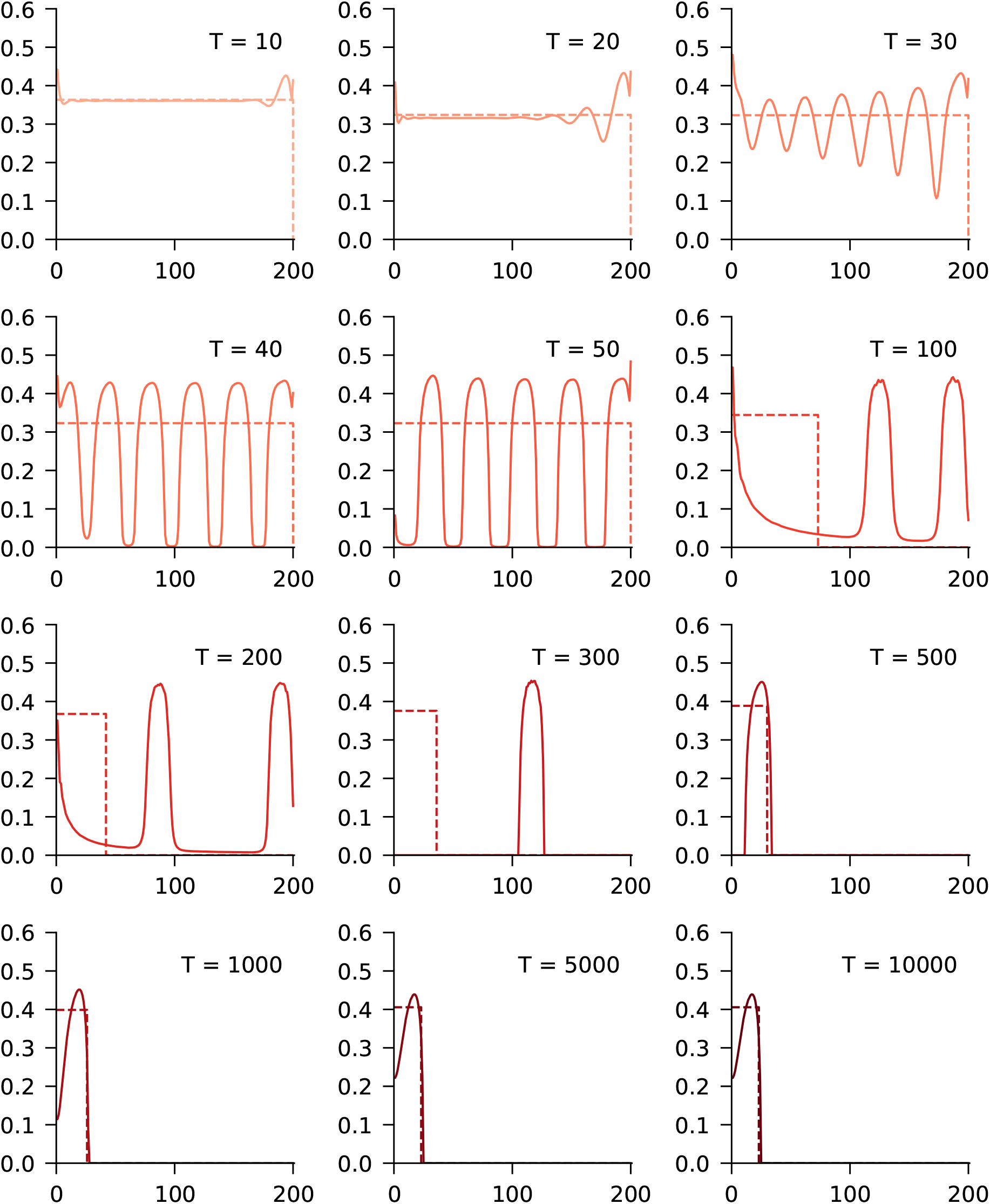
Optimal release patterns for *Wolbachia*. Solid lines are optimal variable release patterns. Dashed lines are optimal constant-density circle release patterns.

**Figure S6.**
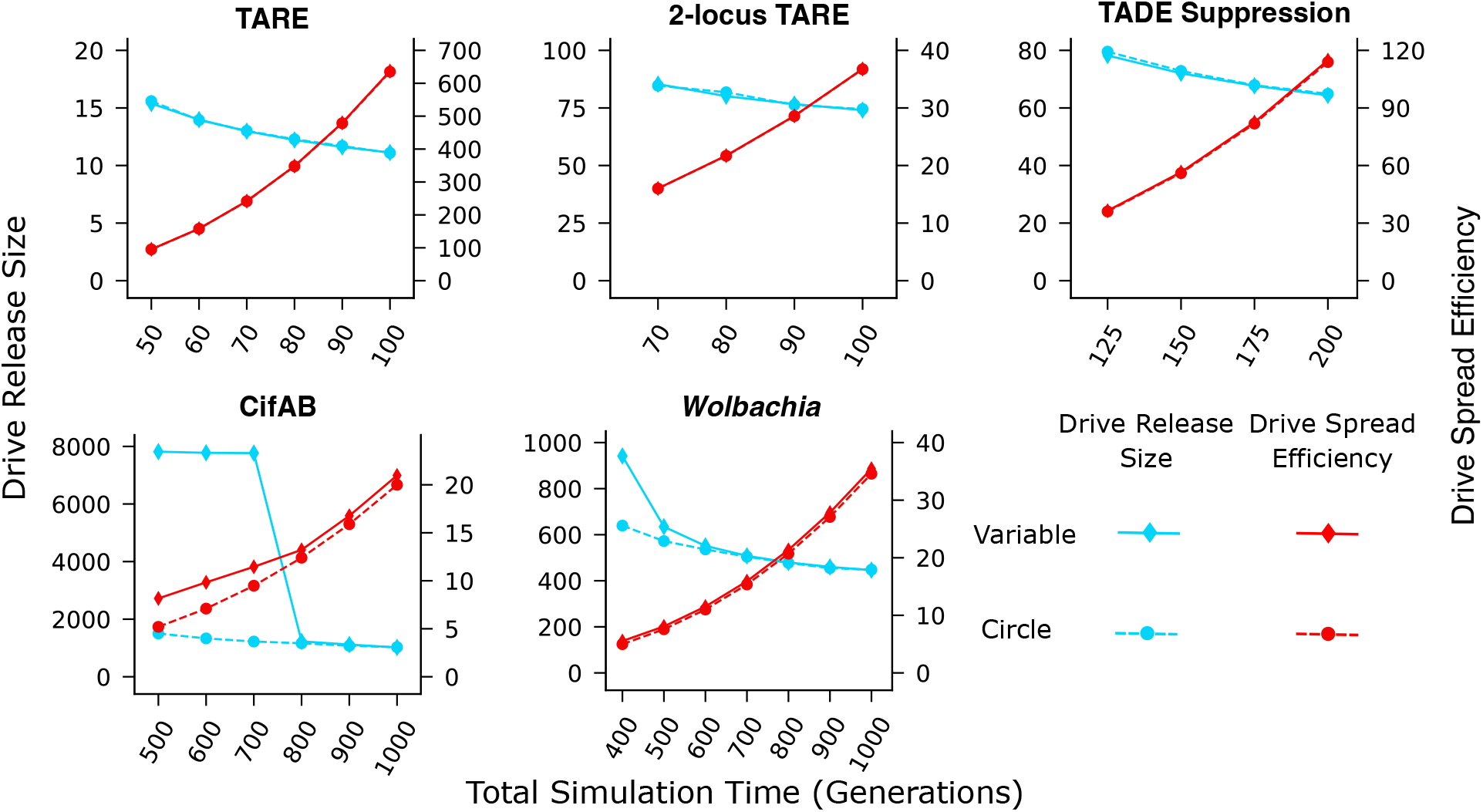
Drive spread efficiency and lower release size. Identical to Figure 6, but with rescaled axis to provide a clearer view of lower release sizes.

**Figure S7.**
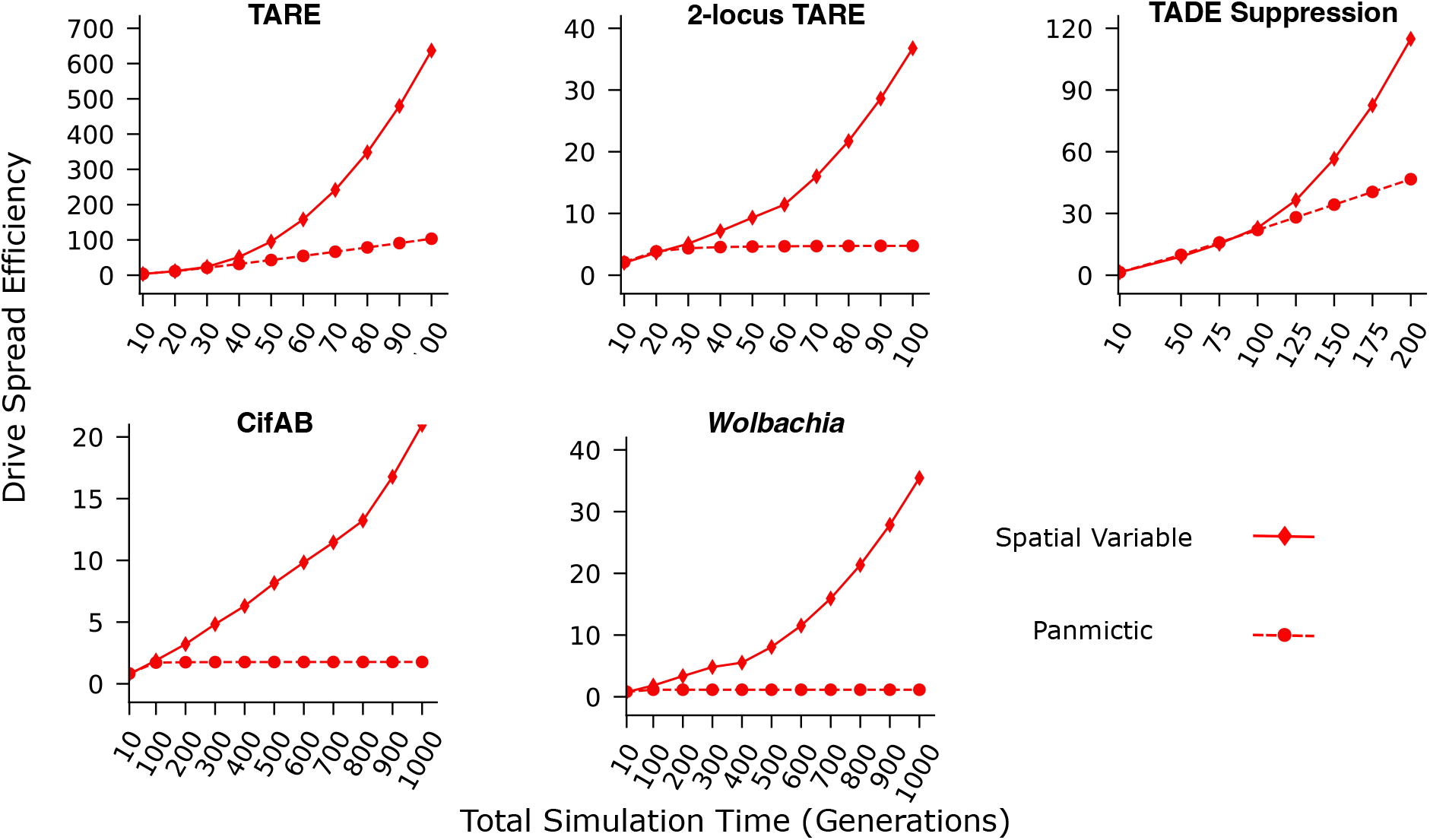
Drive spread efficiency for spatial variable and panmictic releases. For panmictic and variable optimal drive release patterns, we show the drive spread efficiency as a function if simulation timeframe.

### Reaction terms in the reaction diffusion system

We list for each gene drive the **p** in the reaction term 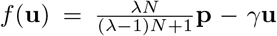. Let *f* denote the frequency of genotype *i* in the local population (i.e., *f*_*i*_ = *u*_*i*_*/N* ). *p*_*i*_ in **p** denotes the probability of getting an offspring of genotype *i* in the next generation. *p*_*i*_’s are not normalized and do not necessarily sum to 1.

#### TARE

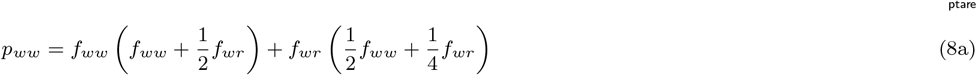

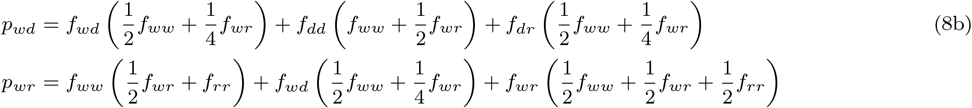

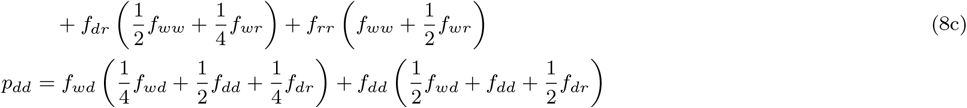

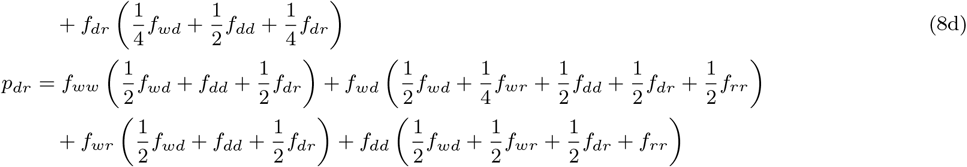

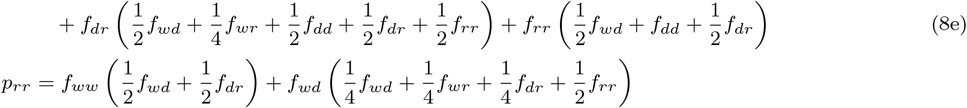

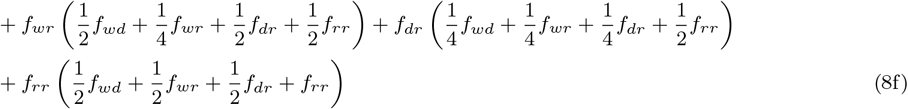

#### 2-Locus TARE

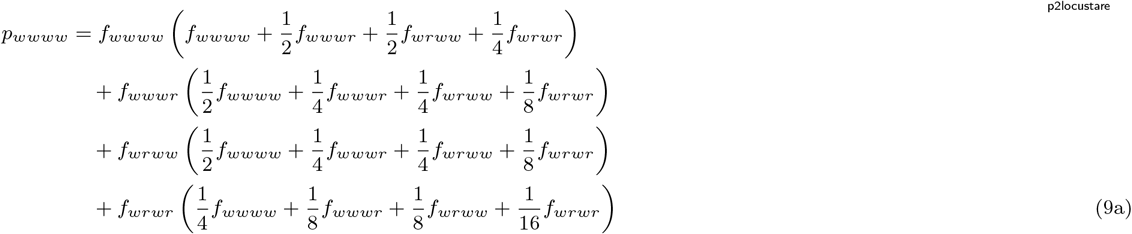

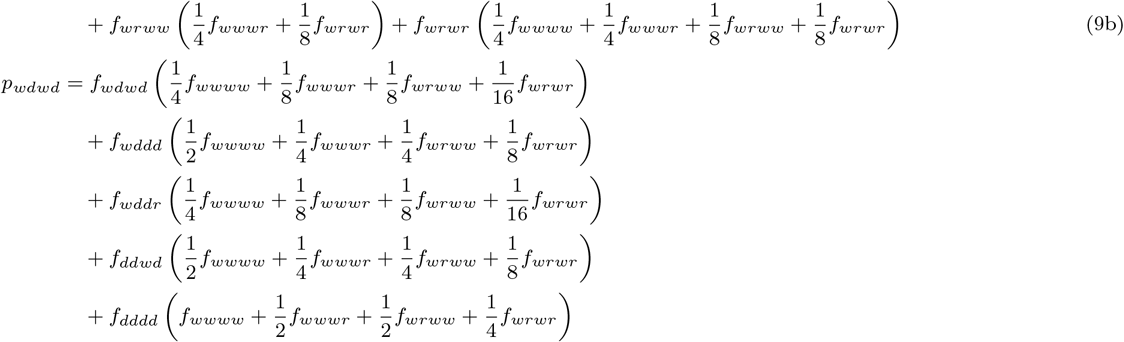

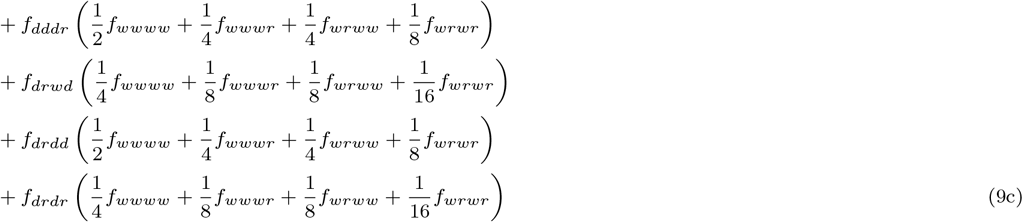

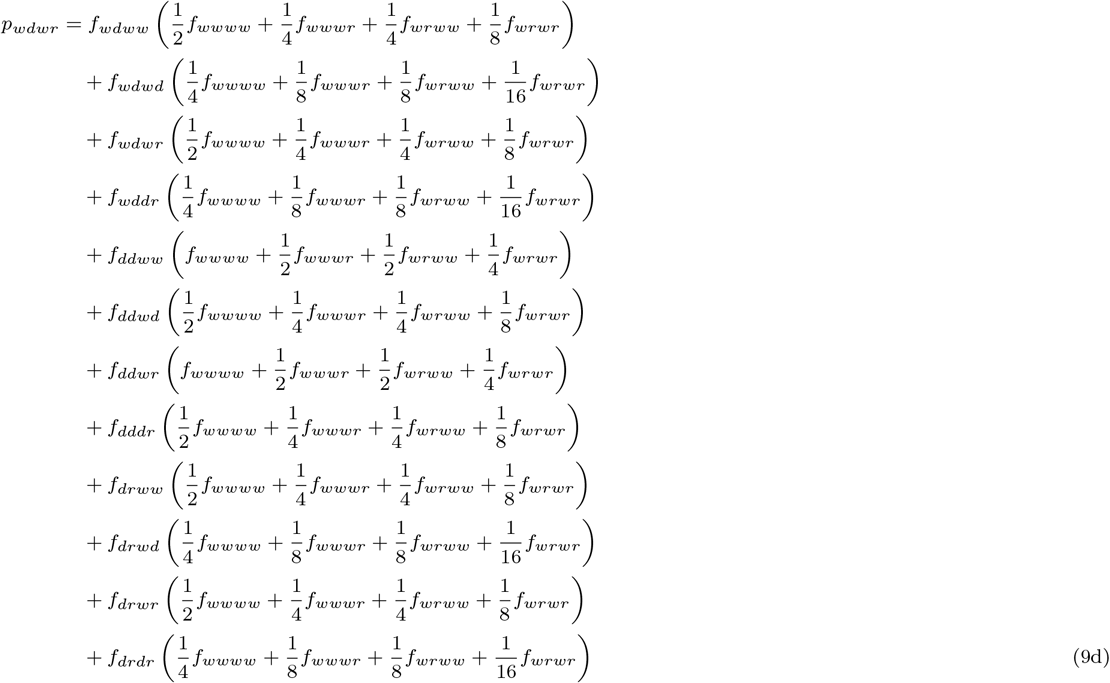

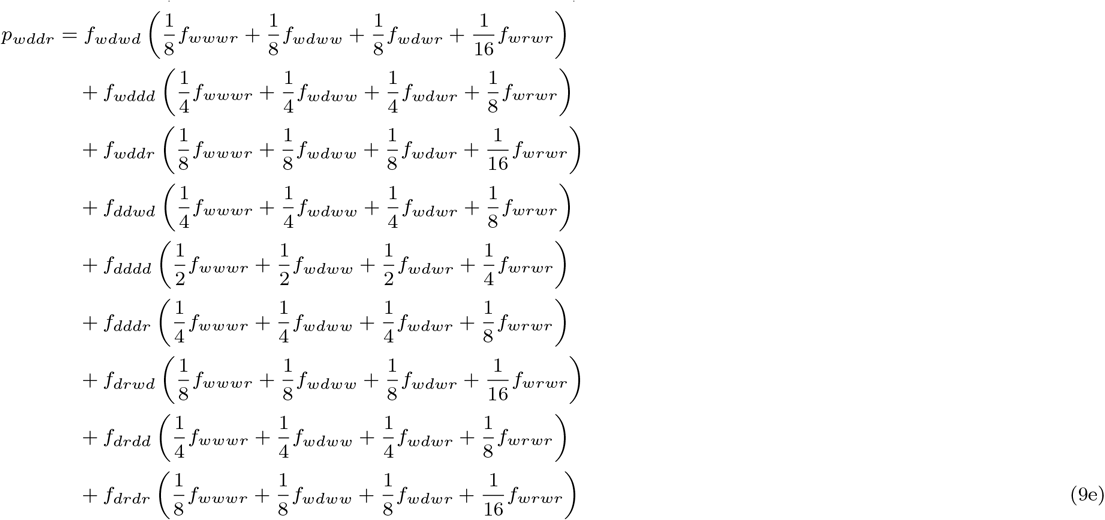

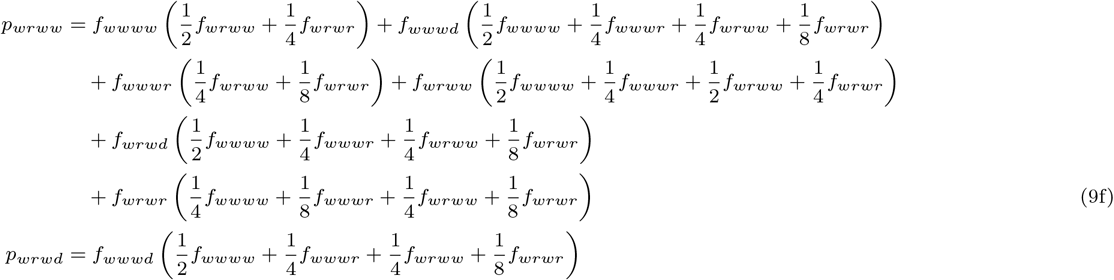

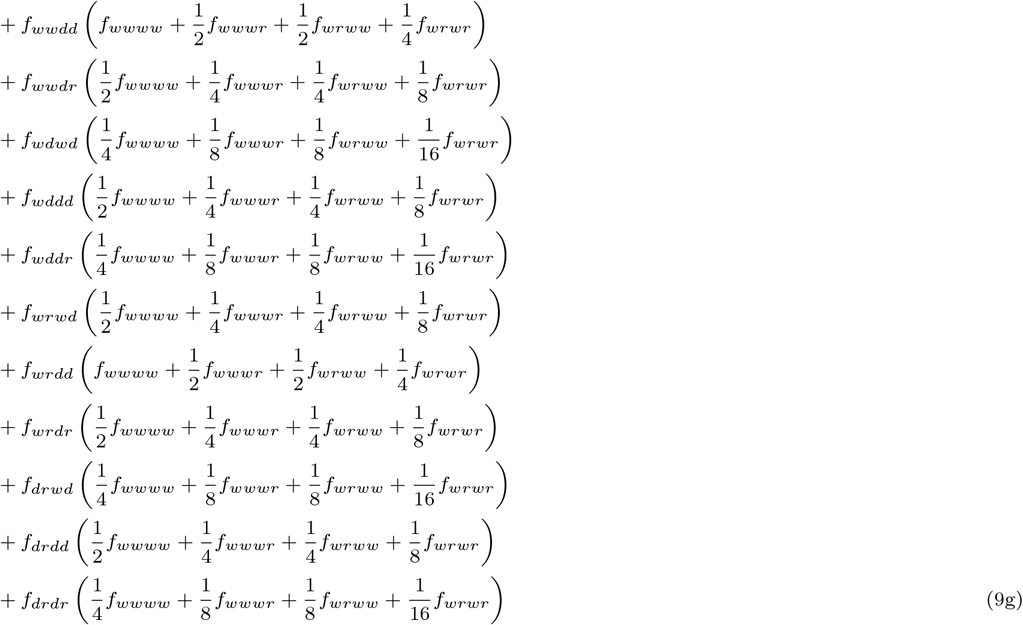

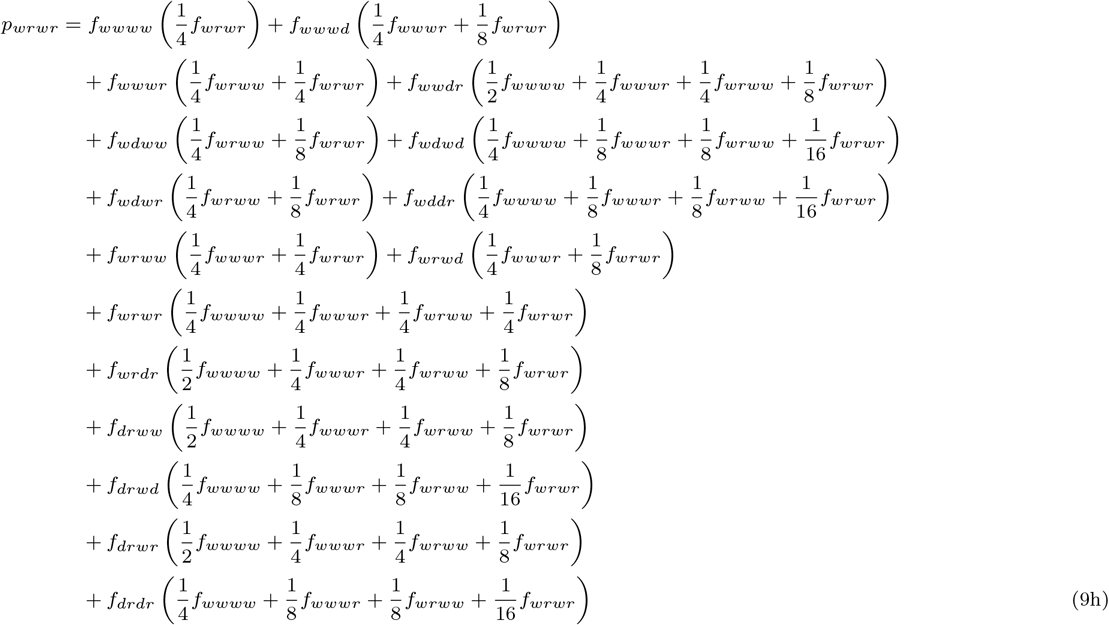

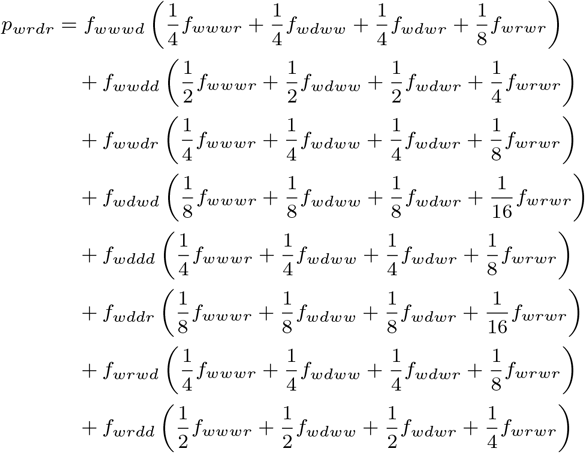

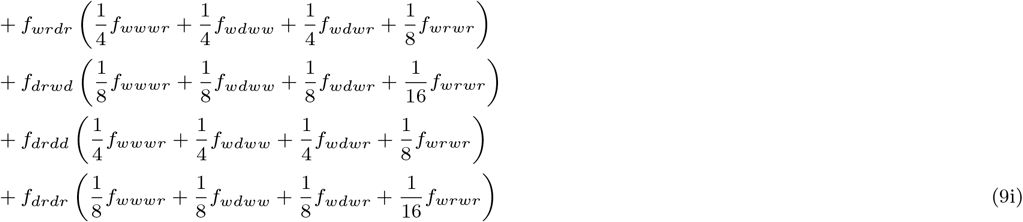

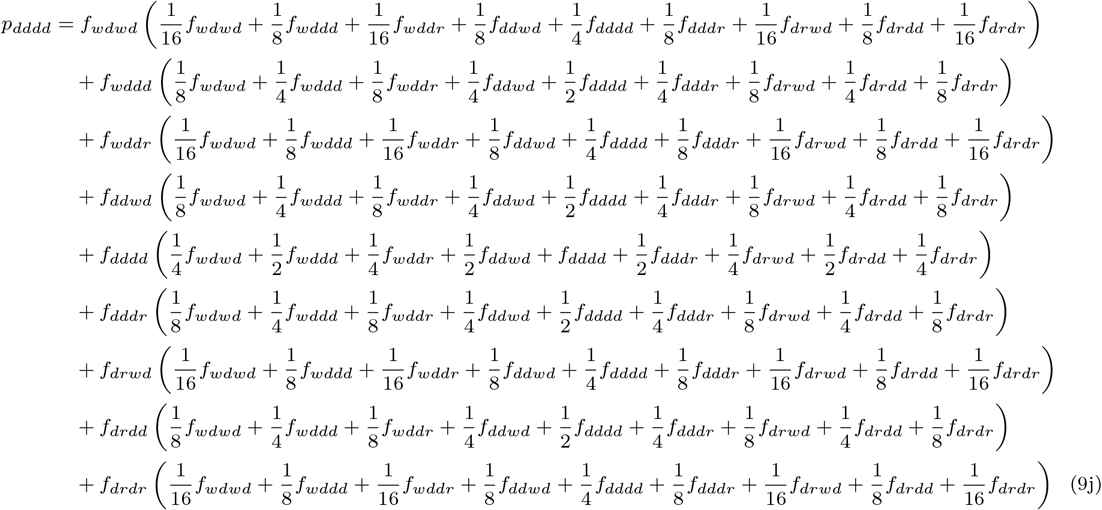

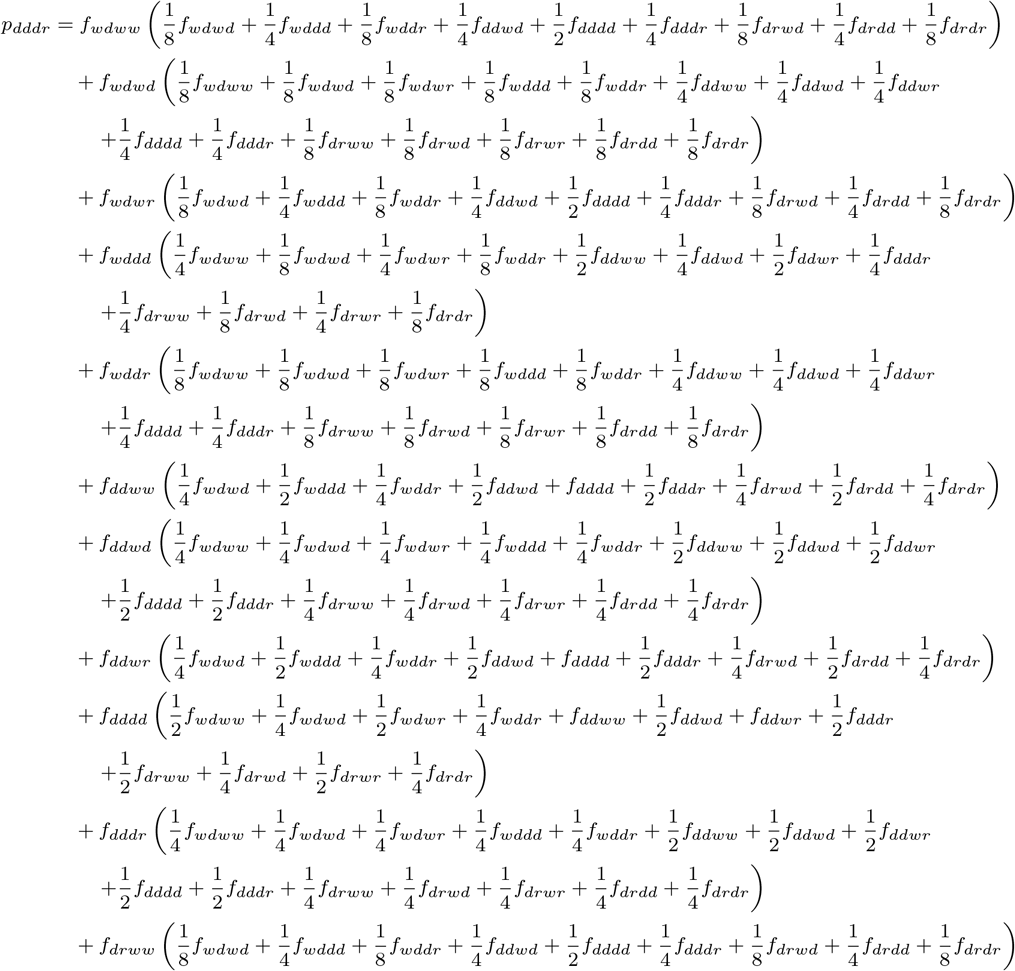

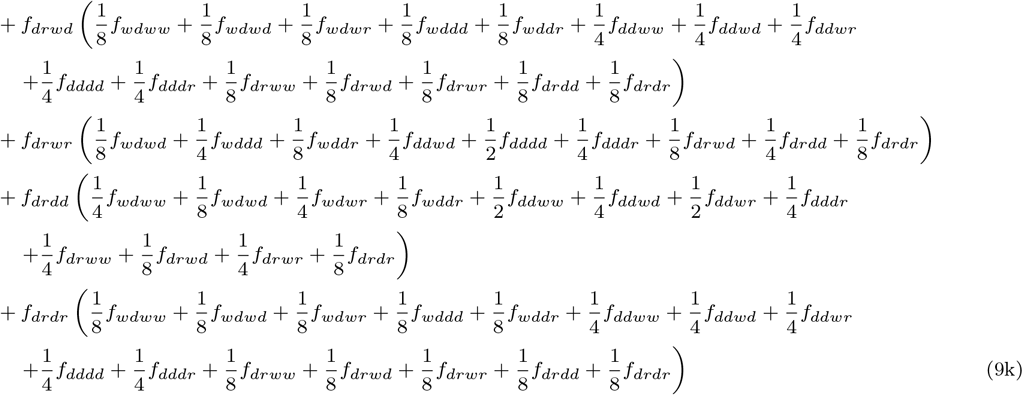

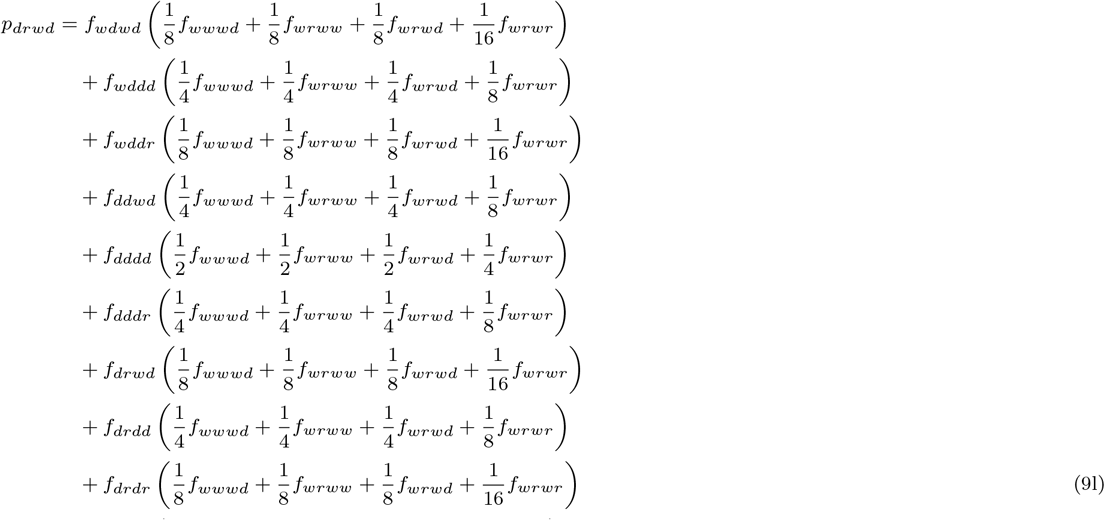

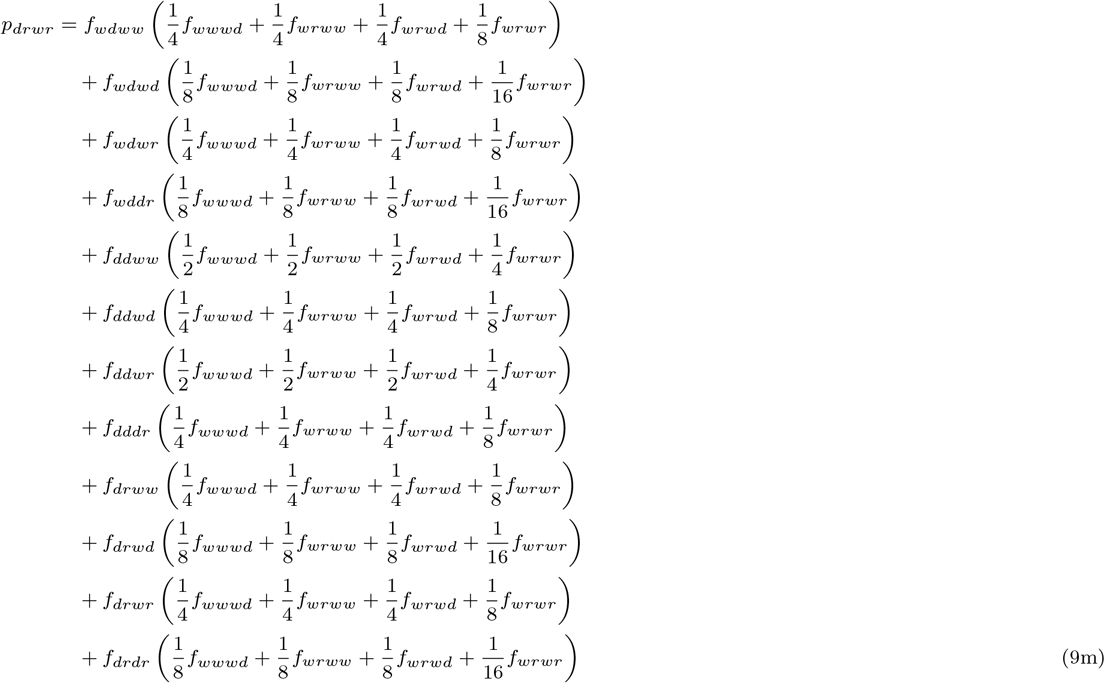

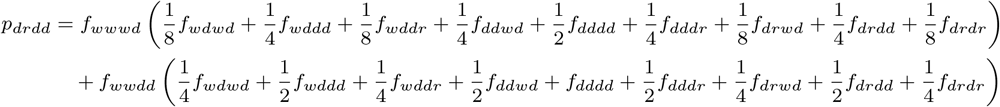

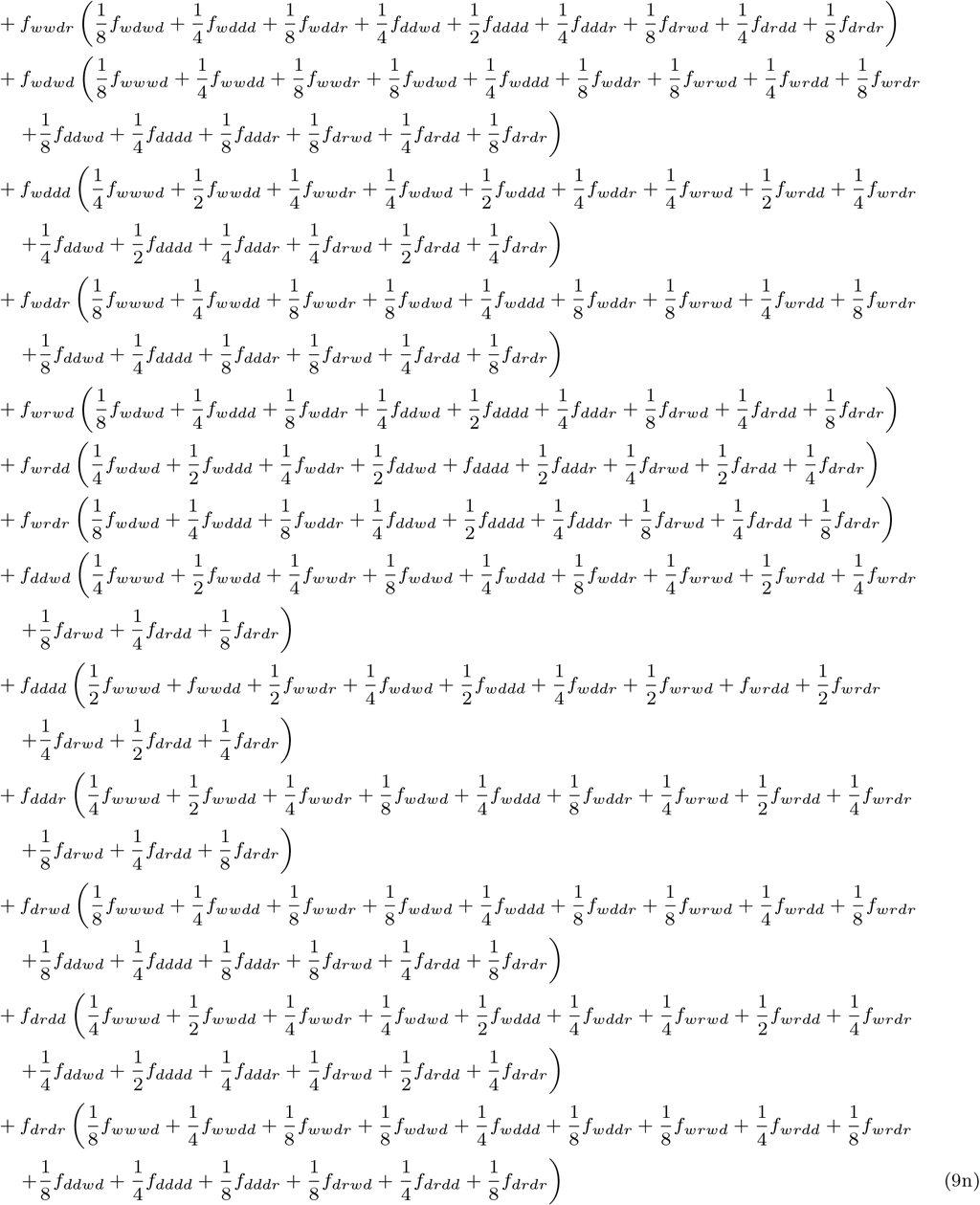

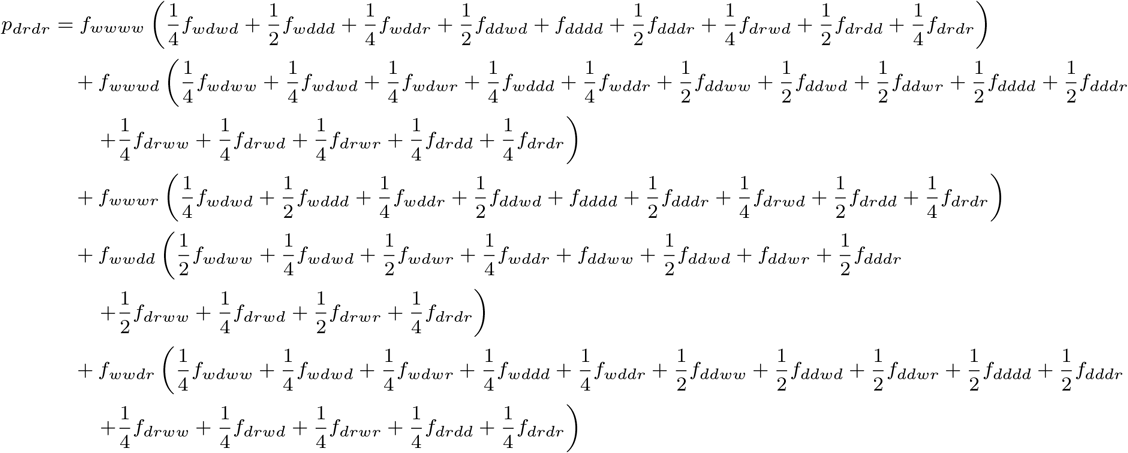

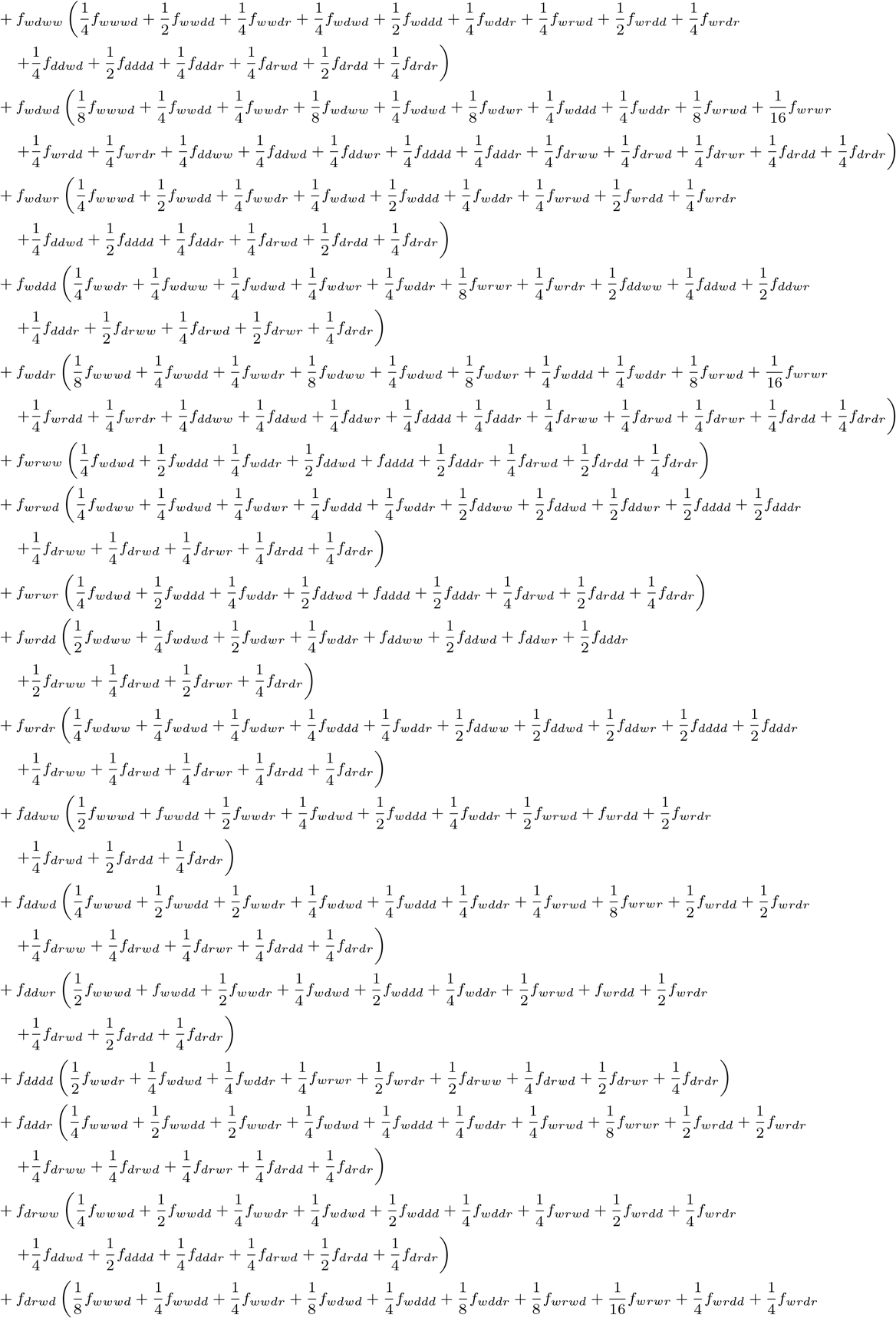

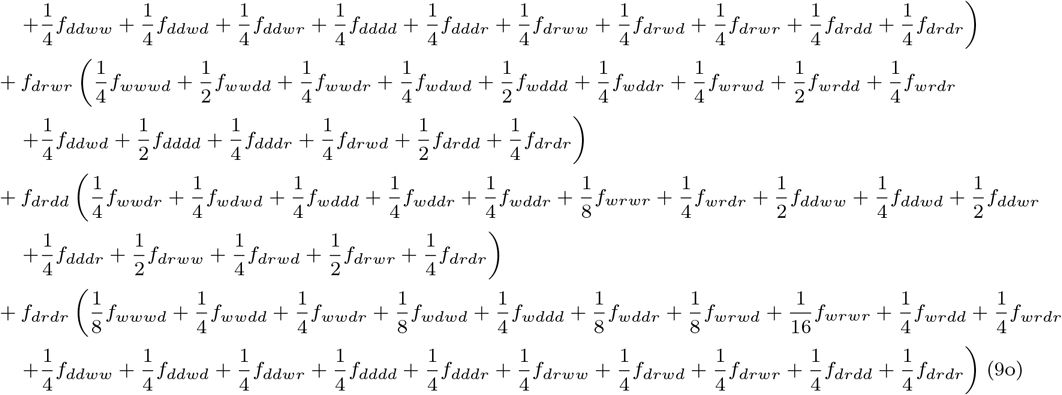

#### TADE Suppression

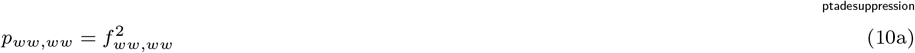

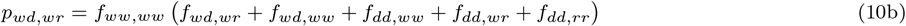

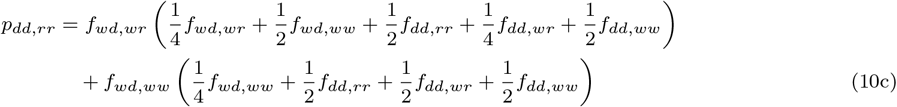

#### CifAB

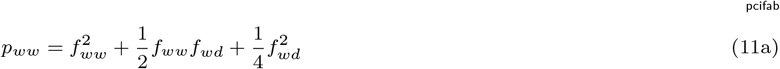

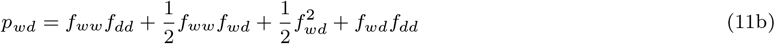

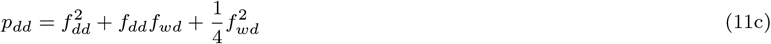

#### Wolbachia

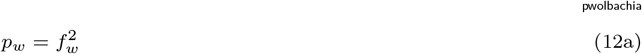

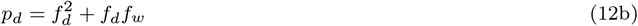

## References

[1] Hay, B. A., Oberhofer, G., and Guo, M. Engineering the composition and fate of wild populations with gene drive. Annu. Rev. Entomol., 66(1):407–434, January 2021.

[2] Verkuijl, S. A. N., Ang, J. X. D., Alphey, L., Bonsall, M. B., and Anderson, M. A. E. The challenges in developing efficient and robust synthetic homing endonuclease gene drives. Front. Bioeng. Biotechnol., 10:856981, March 2022.

[3] Wang, G.-H., Du, J., Chu, C. Y., Madhav, M., Hughes, G. L., and Champer, J. Symbionts and gene drive: two strategies to combat vector-borne disease. Trends Genet., 38(7):708–723, July 2022.

[4] Wang, G.-H., Hoffmann, A., and Champer, J. Gene drive and symbiont technologies for control of mosquito-borne diseases. Annu. Rev. Entomol., 70(1):229–249, January 2025.

[5] Champer, J., Buchman, A., and Akbari, O. S. Cheating evolution: engineering gene drives to manipulate the fate of wild populations. Nat. Rev. Genet., 17(3):146–159, March 2016.

[6] Adelman, Z. N. and Kojin, B. B. Malaria-resistant mosquitoes (diptera: Culicidae); the principle is proven, but will the effectors be effective? J. Med. Entomol., 58(5):1997–2005, September 2021.

[7] Bier, E. Gene drives gaining speed. Nat. Rev. Genet., 23(1):5–22, January 2022.

[8] Leftwich, P. T., Edgington, M. P., Harvey-Samuel, T., Carabajal Paladino, L. Z., Norman, V. C., and Alphey, L. Recent advances in threshold-dependent gene drives for mosquitoes. Biochem. Soc. Trans., 46(5):1203–1212, October 2018.

[9] Dhole, S., Lloyd, A. L., and Gould, F. Gene drive dynamics in natural populations: The importance of density dependence, space, and sex. Annu. Rev. Ecol. Evol. Syst., 51(1):505–531, November 2020.

[10] Champer, J., Yang, E., Lee, E., Liu, J., Clark, A. G., and Messer, P. W. A CRISPR homing gene drive targeting a haplolethal gene removes resistance alleles and successfully spreads through a cage population. Proc. Natl. Acad. Sci. U. S. A., 117(39):24377–24383, September 2020.

[11] Kyrou, K., Hammond, A. M., Galizi, R., Kranjc, N., Burt, A., Beaghton, A. K., Nolan, T., and Crisanti, A. A CRISPR-Cas9 gene drive targeting doublesex causes complete population suppression in caged anopheles gambiae mosquitoes. Nat. Biotechnol., 36(11):1062–1066, December 2018.

[12] Khamis, D., El Mouden, C., Kura, K., and Bonsall, M. B. Ecological effects on underdominance threshold drives for vector control. J. Theor. Biol., 456:1–15, November 2018.

[13] Dhole, S., Vella, M. R., Lloyd, A. L., and Gould, F. Invasion and migration of spatially self-limiting gene drives: A comparative analysis. Evol. Appl., 11(5):794–808, June 2018.

[14] Edgington, M. P. and Alphey, L. S. Conditions for success of engineered underdominance gene drive systems. J. Theor. Biol., 430:128–140, October 2017.

[15] Láruson, J. and Reed, F. A. Stability of underdominant genetic polymorphisms in population networks. J. Theor. Biol., 390:156–163, February 2016.

[16] Huang, Y., Lloyd, A. L., Legros, M., and Gould, F. Gene-drive into insect populations with age and spatial structure: a theoretical assessment: Theoretical assessment of gene-drive. Evol. Appl., 4(3):415–428, May 2011.

[17] Davis, S., Bax, N., and Grewe, P. Engineered underdominance allows efficient and economical introgression of traits into pest populations. J. Theor. Biol., 212(1):83–98, September 2001.

[18] Magori, K. and Gould, F. Genetically engineered underdominance for manipulation of pest populations: A deterministic model. Genetics, 172(4):2613–2620, April 2006.

[19] Altrock, P. M., Traulsen, A., Reeves, R. G., and Reed, F. A. Using underdominance to bi-stably transform local populations. J. Theor. Biol., 267(1):62–75, November 2010.

[20] Edgington, M. P. and Alphey, L. S. Population dynamics of engineered underdominance and killer-rescue gene drives in the control of disease vectors. PLoS Comput. Biol., 14(3):e1006059, March 2018.

[21] Champer, J., Zhao, J., Champer, S. E., Liu, J., and Messer, P. W. Population dynamics of under-dominance gene drive systems in continuous space. ACS Synth. Biol., 9(4):779–792, April 2020.

[22] Champer, J., Champer, S. E., Kim, I. K., Clark, A. G., and Messer, P. W. Design and analysis of CRISPR-based underdominance toxin-antidote gene drives. Evol. Appl., 14(4):1052–1069, April 2021.

[23] Chen, C.-H., Huang, H., Ward, C. M., Su, J. T., Schaeffer, L. V., Guo, M., and Hay, B. A. A synthetic maternal-effect selfish genetic element drives population replacement in drosophila. Science, 316 (5824):597–600, April 2007.

[24] Champer, J., Lee, E., Yang, E., Liu, C., Clark, A. G., and Messer, P. W. A toxin-antidote CRISPR gene drive system for regional population modification. Nat. Commun., 11(1):1082, February 2020.

[25] Champer, J., Kim, I. K., Champer, S. E., Clark, A. G., and Messer, P. W. Performance analysis of novel toxin-antidote CRISPR gene drive systems. BMC Biol, 18(1), December 2020.

[26] Oberhofer, G., Ivy, T., and Hay, B. A. Cleave and rescue, a novel selfish genetic element and general strategy for gene drive. Proc. Natl. Acad. Sci. U. S. A., 116(13):6250–6259, March 2019.

[27] Oberhofer, G., Ivy, T., and Hay, B. A. Gene drive and resilience through renewal with next generation cleave and rescue selfish genetic elements. Proc. Natl. Acad. Sci. U. S. A., 117(16):9013–9021, April 2020.

[28] Feng, R. and Champer, J. Deployment of tethered gene drive for confined suppression in continuous space requires avoiding drive wave interference. Mol. Ecol., 33(19):e17530, 2024.

[29] Metzloff, M., Yang, E., Dhole, S., Clark, A. G., Messer, P. W., and Champer, J. Experimental demonstration of tethered gene drive systems for confined population modification or suppression. BMC Biol., 20(1):119, May 2022.

[30] Dhole, S., Lloyd, A. L., and Gould, F. Tethered homing gene drives: A new design for spatially restricted population replacement and suppression. Evol. Appl., 12(8):1688–1702, 2019.

[31] Schmidt, T. L., Filipović, I., Hoffmann, A. A., and Rašić, G. Fine-scale landscape genomics helps explain the slow spatial spread of wolbachia through the aedes aegypti population in cairns, australia. Heredity (Edinb.), 120(5):386–395, 2018.

[32] Schmidt, T. L., Barton, N. H., Rašić, G., Turley, A. P., Montgomery, B. L., Iturbe-Ormaetxe, I., Cook, P. E., Ryan, P. A., Ritchie, S. A., Hoffmann, A. A., O’Neill, S. L., and Turelli, M. Local introduction and heterogeneous spatial spread of dengue-suppressing wolbachia through an urban population of aedes aegypti. PLoS Biol., 15(5):e2001894, May 2017.

[33] Fisher, R. A. The wave of advance of advantageous genes. Ann. Eugen., 7(4):355–369, June 1937.

[34] Barton, N. H. The dynamics of hybrid zones. Heredity (Edinb.), 43(3):341–359, December 1979.

[35] Piálek, J. and Barton, N. H. The spread of an advantageous allele across a barrier: the effects of random drift and selection against heterozygotes. Genetics, 145(2):493–504, February 1997.

[36] Barton, N. H. and Turelli, M. Spatial waves of advance with bistable dynamics: cytoplasmic and genetic analogues of allee effects. Am. Nat., 178(3):E48–75, September 2011.

[37] Rouhani, S. and Barton, N. Speciation and the “shifting balance” in a continuous population. Theor. Popul. Biol., 31(3):465–492, June 1987.

[38] Li, J. and Champer, J. Harnessing wolbachia cytoplasmic incompatibility alleles for confined gene drive: A modeling study. PLoS Genet., 19(1):e1010591, January 2023.

[39] Florez, D., Cortez, R., Hyman, J. M., and Qu, Z. Improving wolbachia-based control programs in urban settings: Insights from spatial modeling. PLoS Negl. Trop. Dis., 19(12):e0013787, 2025.

[40] Kim, I. K. and Messer, P. W. Predicting the invasiveness of threshold-dependent gene drives. bioRxivorg, page 2025.11.20.689598, 2025.

[41] Pan, M. and Champer, J. Making waves: Comparative analysis of gene drive spread characteristics in a continuous space model. Mol. Ecol., 32(20):5673–5694, October 2023.

[42] Zhu, Y. and Champer, J. Simulations reveal high efficiency and confinement of a population suppression CRISPR toxin-antidote gene drive. ACS Synth. Biol., 12(3):809–819, March 2023.

[43] Zhang, S. and Champer, J. Performance characteristics allow for confinement of a CRISPR toxin-antidote gene drive for population suppression in a reaction-diffusion model. Proc. Biol. Sci., 291 (2025):20240500, June 2024.

[44] LePage, D. P., Metcalf, J. A., Bordenstein, S. R., On, J., Perlmutter, J. I., Shropshire, J. D., Layton, E. M., Funkhouser-Jones, L. J., Beckmann, J. F., and Bordenstein, S. R. Prophage WO genes recapitulate and enhance wolbachia-induced cytoplasmic incompatibility. Nature, 543(7644): 243–247, March 2017.

[45] Shropshire, J. D. and Bordenstein, S. R. Two-by-one model of cytoplasmic incompatibility: Synthetic recapitulation by transgenic expression of cifA and cifB in drosophila. PLoS Genet., 15(6):e1008221, June 2019.

[46] Adams, K. L., Abernathy, D. G., Willett, B. C., Selland, E. K., Itoe, M. A., and Catteruccia, F. Wolbachia cifB induces cytoplasmic incompatibility in the malaria mosquito vector. Nat. Microbiol., 6(12):1575–1582, December 2021.

[47] Hancock, P. A., Ritchie, S. A., Koenraadt, C. J. M., Scott, T. W., Hoffmann, A. A., and Godfray, H. C. J. Predicting the spatial dynamics of Wolbachia infections in Aedes aegypti arbovirus vector populations in heterogeneous landscapes. J. Appl. Ecol., 56(7):1674–1686, 2019.

[48] Beaghton, A., Hammond, A., Nolan, T., Crisanti, A., Godfray, H. C. J., and Burt, A. Requirements for driving antipathogen effector genes into populations of disease vectors by homing. Genetics, 205 (4):1587–1596, April 2017.

[49] Turelli, M. and Barton, N. H. Deploying dengue-suppressing wolbachia : Robust models predict slow but effective spatial spread in aedes aegypti. Theor. Popul. Biol., 115:45–60, 2017.

[50] Bull, J. J., Remien, C. H., and Krone, S. M. Gene-drive-mediated extinction is thwarted by population structure and evolution of sib mating. Evol. Med. Public Health, 2019(1):66–81, 2019.

[51] Zhang, X., Sun, W., Kim, I. K., Messer, P. W., and Champer, J. Population dynamics in spatial suppression gene drive models and the effect of resistance, density dependence, and life history. bioRxivorg, page 2024.08.14.607913, 2024.

